# Differential Assembly of Mouse and Human Tumor Microenvironments

**DOI:** 10.1101/2025.07.01.661217

**Authors:** Tristan Courau, Rebecca G. Jaszczak, Bushra Samad, Emily Flynn, Nayvin W. Chew, Gabriella C. Reeder, Jessica Tsui, Arja Ray, Harrison Wismer, Daniel Bunis, Leonard Lupin-Jimenez, Noah V. Gavil, David Masopust, John P. Graham, Daniel A. Skelly, Xavier Vesco, Edison T. Liu, Gabriela K. Fragiadakis, Alexis J. Combes, Matthew F. Krummel

## Abstract

Mouse models are frequently used to develop treatments for human cancer. Yet, we lack a comprehensive understanding of the comparative organization of mouse and human tumor microenvironments (mu/huTMEs). Through immunoprofiling of commonly used mouse models, we found that the immune composition of most muTMEs resemble poorly infiltrated human tumors extensively biased toward high macrophages densities. Relatedly, we discover species-specific biases of chemokine expression networks, factors which drive TMEs assembly. Further, assessing coarse cellular networks, we find conserved correlations between some immune cell frequencies, while other relationships only appear conserved in the huTMEs matching muTME profiles. Despite this variable alignment, we define robust cell type-specific gene expression programs conserved in TMEs across species and cohorts and identify ones that are coordinated between cell populations in both species. Together, we isolate and offer methods to study the multiple areas of hazard and opportunities for using mice to model human cancer.

**Graphical Abstract:** 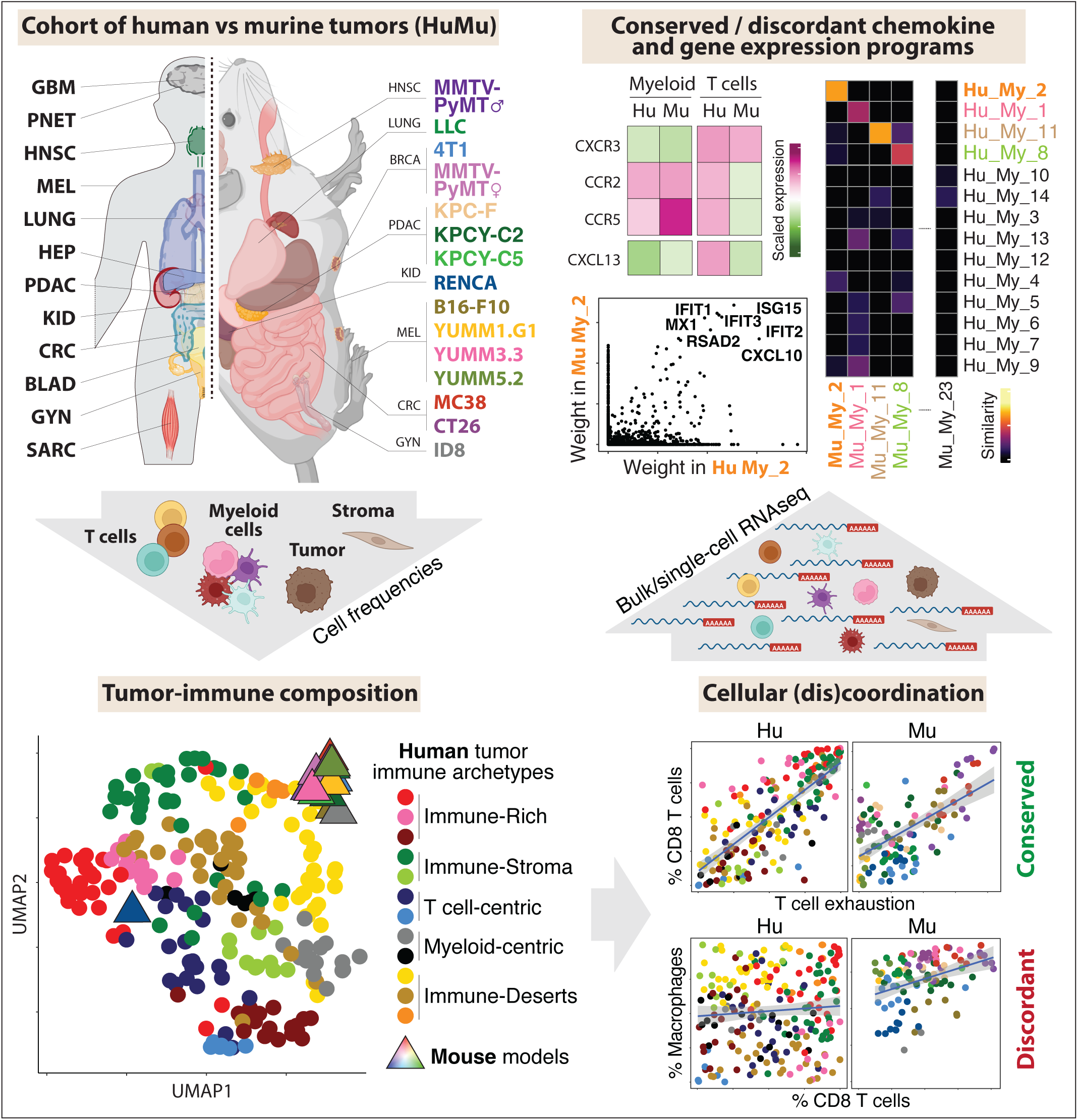

## Introduction

The immune composition of human tumor microenvironments (huTMEs) is now established as a crucial factor for patients outcome^1,2^. There is a great diversity of huTMEs, ranging from a number of distinct subsets that are highly inflamed and immune-rich (collectively called “hot”) to at least three types that are poorly infiltrated (collectively, “cold”), each subdivided by their relative amounts of fibrotic components as well as T cells and myeloid composition^3–7^. These classes of huTMEs are associated with disease outcomes across various cancer tissues of origin^3–7^ and each likely require variations in drug design to generate cures. However, the biological mechanisms underlying the formation of these recurring TME patterns remain poorly understood. This represents a critical need for designing treatment decisions based on their specific features and sensitivities^8,9^.

Despite millions of years of evolutionary divergence between mouse and human, mice remain invaluable for studying human biology. When investigating malignant cell biology, mouse models have historically provided critical insights into conserved mechanisms of tumor development, progression, and drug resistance^10^. Immunologically, mice have been notable for the original demonstration of checkpoint blockades, as well as for elucidating and targeting key immune escape mechanisms used by tumors^11–13^. Despite their major importance in the development of successful treatments^14,15^, numerous failures in predictive efficacy of mouse modeling^16,17^ suggests a need for both more nuanced model selection and more detailed understanding of the limitations of mouse models to study human cancers.

In-depth comparison of TMEs between these species has been limited. A small number of studies have compared the murine TMEs (muTMEs) of a few selected mouse models, overall concluding the presence of some degree of diversity in their immune profiles, noting also that some vary with tumor size^18–22^. Other groups compared the TMEs of a small number of mouse models to indication-matched patients, and find discrepancies in their immunogenicity and T cell infiltration profiles^23^. While insightful, these studies did not provide enough resolution, nor did they consider the diversity of huTMEs diversity outside of a single indication space, to assess the common relevance of mouse tumor models in substantial detail.

A few frameworks currently conceptualize the diversity of TMEs found in cancer patients. Holistic categorizations of the cells and their gene expression has led to discovery of conserved TMEs “subtypes”^3,4^, “ecotypes”^6^ or “archetypes”^7^ which place huTMEs into classes and are presumed to represent a form of template for local immune systems. In such data, some archetypes most closely resemble non-healing wounds^9,24^ and have co-opted this biology whereas others may, for example, more closely resemble the immune détente surrounding chronic viral infection^9,25–29^. These holistic classifications often contain some elements of the immune ‘rich’ vs. ‘poor’ designations, but with significantly greater granularity. Another approach to compare immune systems across species and TMEs is to characterize the intimate cell-cell relationships between cell types as a building block of immune function. For example, the frequencies of Treg and exhausted T cells have each been linked to the frequencies of certain myeloid populations^30–33^, some but not all of which have been found to correlate also in some huTMEs. In another example, some tumors were found to have significant B cell content, often in spatial aggregates and correlated with dendritic cells and various stromal (fibroblast) populations, together described as tertiary lymphoid structures (TLS)^34^. In human and mice, the occurrence of TLS—sometimes called out as cellular ‘neighborhoods’ using imaging methods—has been found to have generally significant positive prognostic value^34^ and in human may be mediated through co-inclusion of CD4 T cells expressing the B cell-attracting chemokine CXCL13^35–37^.

To move towards a systematic comparison of TMEs across species and understand fundamental similarities and differences that affect study interpretation, we collected high dimensional data across a variety of muTMEs, and then systematically compared holistic views (e.g. archetypes/immune subtypes) as well as gene-expression programs (GEPs) with existing huTMEs. The data compendium thus assembled allowed us to discover typical and penetrant biases in specific chemokine expression (including CXCL13), in cell-cell relationships and in the conservation and/or co-occurrence of robust GEPs in both species. Altogether, this resource provides rich data as well as a first set of critical analyses of the TMEs as they represent across species. These in turn will allow us and others to judiciously select and support studies of mouse tumors as models for human disease, to seek specific features in each model, and to provide milestones for refining them and their typical usage.

## Results

### Most mouse models only recapitulate the immune composition of low-infiltration, macrophage-rich human tumors

In an attempt to benchmark the immune composition and transcriptomic patterns of typical muTMEs relative to huTMEs, we first collected high-dimensional data on 15 murine models, selecting the most widely used ones in the field, estimated to represent >95% of all published immunotherapy studies. This selection includes commonly used cell lines such as B16F10, MC38, CT26, LLC, and 4T1, as well as a selection of models that are autochthonous/transplanted (KPC) and genetically engineered (MMTV-PyMT), and in the two most commonly used background strains of mice (BALB/c and C57BL/6)^18–20,23,38^ (**Figure 1A**). While we recognize that other models and variables may be missing from our selection, the size of datasets as well as the desire to answer the broad question in the most widely used systems provided the basis for picking a representative 15 mouse models for our study. To ensure consistency, tumors formed from these different mouse models were analyzed at day 14 post tumor implantation (or ∼500mm^3^). All were analyzed by mass cytometry (CyTOF) for compositional analysis and 9 of them were additionally processed for single-cell RNA sequencing (scRNAseq) to discover gene expression programs.

**Figure 1:**
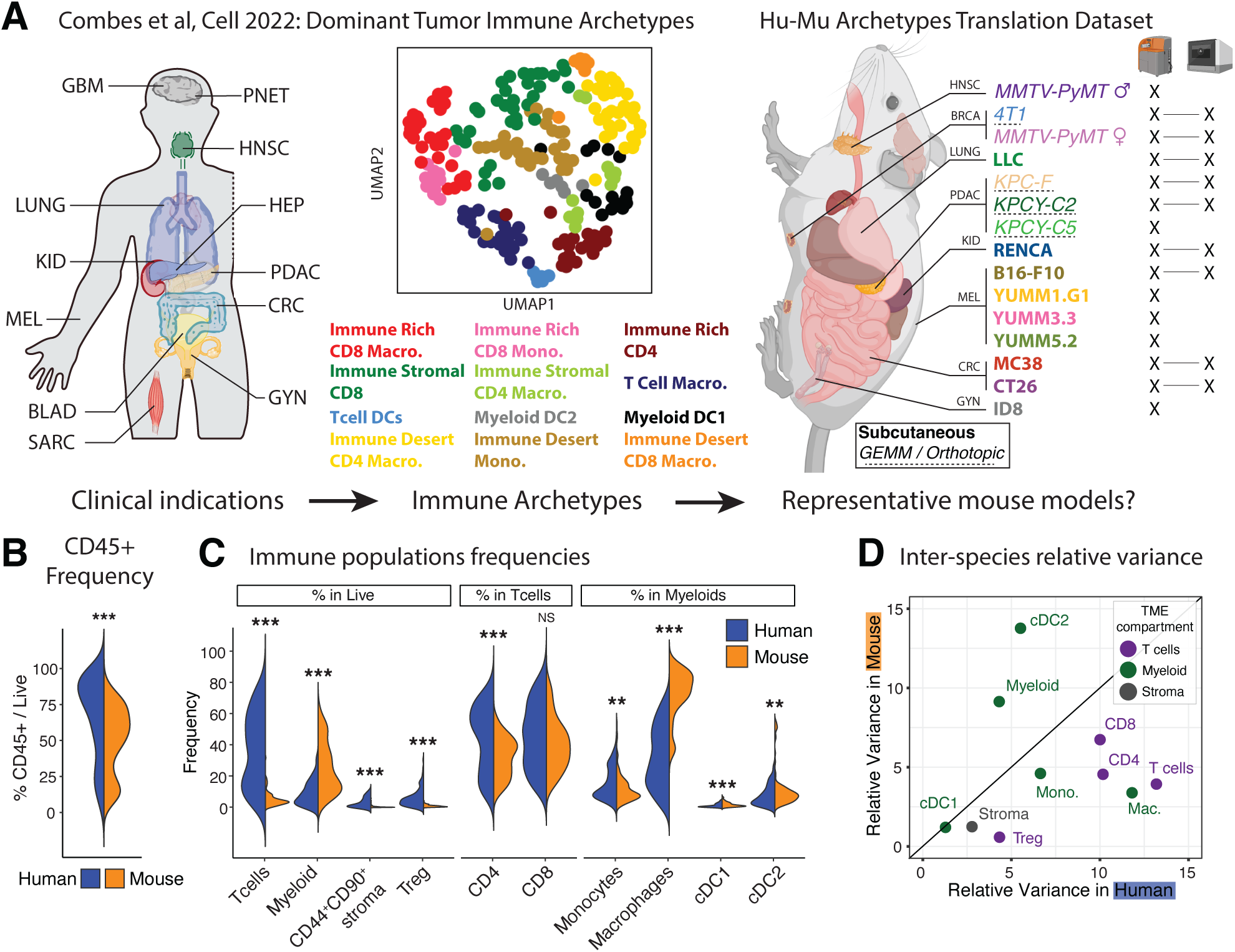
Compositional disparities between human and murine tumors. **A.** Schematic of our human and murine study cohorts. **B.** Violin plot presenting the frequency of CD45+ in Live cells from human (blue, all samples grouped, n = 224) vs murine (orange, all samples grouped, n = 109) tumors. **C.** Violin plot presenting the frequencies of T cells (Tregs excluded), non-granulocytic myeloid cells (combining monocytes, macrophages, conventional DCs and plasmacytoid DCs), non-immune stroma (CD44+CD90+ in CD45-) and CD4+ Tregs out of Live cells; CD4 T conventional and CD8 out of T cells; Monocytes, Macrophages, cDC1 and cDC2 out of myeloid cells between human and mouse tumors. **D.** Plot comparing the relative variance of each parameter shown in B. (colored by cellular compartment) between human and murine tumors. The diagonal line represents an equal relative variance between the 2 species. Statistical significance was calculated using a t-test with Bonferroni correction, * p.adj ≤ 0.05, ** p.adj ≤ 0.01, *** p.adj ≤ 0.001.

As a primary comparator, we used the human ImmunoProfiler dataset for which dominant immune archetypes were previously defined based on compositional clustering of human tumors^7^. That dataset similarly defined immune composition while capturing deep RNA sequencing of T cell, non-granulocytic myeloid, stromal and tumor populations and has benchmarked multiple studies and meta-studies^7,31,39,40^. Reasoning that the nature of immune systems are at least partially functions of the density of total or individual cell types, we first analyzed the frequencies of total immune cells across huTMEs and muTMEs (**Figure S1A**). This revealed that while the distribution of TMEs in both species have a pronounced bimodal distribution of total immune density (representing ‘rich’ and ‘poor’ tumors), the global frequencies of immune cells in muTMEs as a fraction of total tissue was significantly lower than the distribution of these frequencies in huTMEs (**Figure 1B**).

Based on our studies and others that examine fundamental cell types to classify huTMEs^4,6,7^, we next focused on the frequencies of 10 major cellular compartments of the TME (**Figure S1A-B**), inspired by the human ImmunoProfiler approach^7^. The first four represent broad compositional features: total effector T cells (including both CD4+ conventional and CD8+ T cells), CD4+ regulatory T cells (Treg), non-granulocytic myeloid cells (simplified as “myeloid” across this manuscript) and CD44^+^CD90^+^ non-immune stromal cells (simplified as “stroma”). The additional six features represent the densities of subsets of T and myeloid cells, namely the fraction of CD4+ and CD8+ conventional T cells (Tconv), and the fractional representation of cells within the “myeloid” compartment, namely tumor-associated macrophages, monocytes and classical dendritic cells type 1 (cDC1) and type 2 (cDC2). Quantifying these densities, we found that muTMEs generally contain significantly lower frequencies of total T cells and higher frequencies of myeloid cells, especially of macrophages, compared to the compendium of huTMEs (**Figure 1C**). That T cells vary much more widely in huTMEs as compared to muTMEs was also seen by relative variance analysis (**Figure 1D**) where comparatively the variance in muTMEs is mostly due to myeloid cells density and composition.

We then sought to use these parameters—which effectively parse human TMEs into archetypes—to see where these commonly used models each parse relative to various human TME biology. Utilizing a UMAP embedding, we first assessed the overall similarity between muTMEs and huTMEs, scaling the 10 previously described features of human and mouse samples together (using z-score) to visualize them in the same lower dimensional space (**Figure 2A**). The UMAP embedding of only muTMEs composition primarily grouped samples by tumor cell line (**Figure S1C-D**), in line with previous reports^18–20,23^. However, embedding both mouse and human samples together revealed that all mouse models, except RENCA, tended to group together with immune-desert, macrophage-rich human samples (**Figure 2A**). To quantify this, we calculated the cosine similarity between human archetypes and mouse models and displayed these similarities as a hierarchically clustered heatmap (**Figure 2B**). This analysis confirmed that RENCA tends to cluster with both immune-stromal CD4-biased (IS CD4) and the two myeloid-centric archetypes (MC DC1 and MC DC2), while all other 14 models studied cluster closely with the immune desert CD4- and CD8-biased, macrophage-rich archetypes (ID CD4/CD8 Mac). Overall, these two archetypes account for only 17% of the patients included in our ImmunoProfiler cohort^7^, with indication-specific variations ranging from 0% in hepatocellular carcinomas to 4.5% in melanoma, 18% in colorectal cancer and 46.5% in gynecological tumors (**Figure S1E)**.

**Figure 2:**
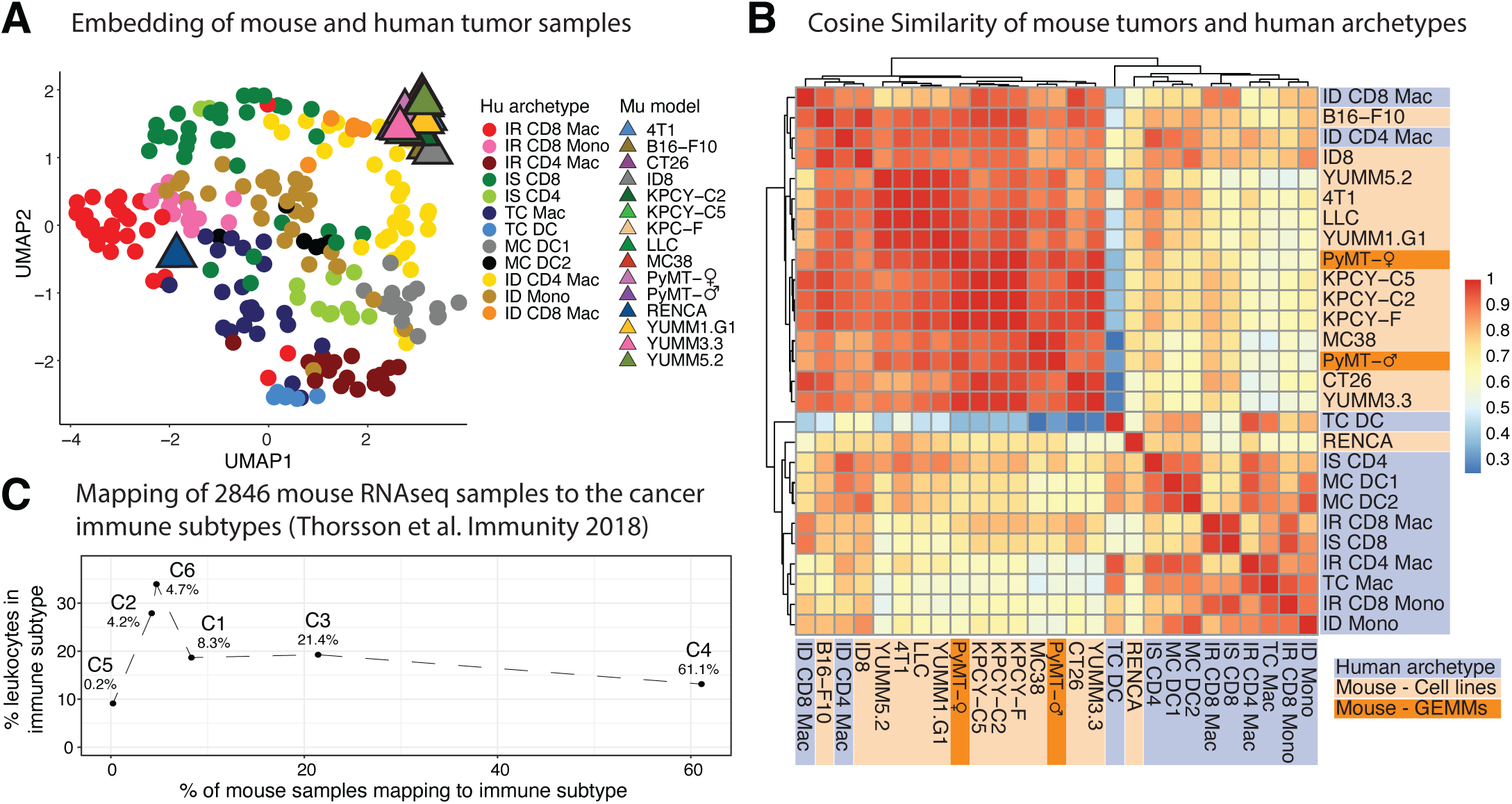
Mapping mouse tumor models to human TME archetypes and subtypes. **A.** Embedding of the frequencies in Figure 1C showing human samples (circles, individual samples colored by archetypes) and mouse tumors (triangles, averaged by tumor line) in the same UMAP space. The frequencies of each population were z-scored across all mouse and human samples before performing UMAP embedding of all 10 z-scored parameters. **B.** Hierarchically clustered matrix of cosine similarities calculated between each mouse tumor model (orange) and each human archetype (blue), using the frequencies in Figure 1C. **C.** Scatter plot presenting the mapping of 2846 mouse tumor samples from NCBI Sequence Read Archive in the 6 Immune Subtypes described in^3^. The graph shows the percentage of mouse samples mapping to each subtype (X axis, values also displayed on the graph) as well as the average percentage of leukocytes infiltrated in each subtype (Y axis).

Since we had only included 15 mouse models in our study, we sought to extend the similarity analysis further and therefore queried RNAseq data from a cohort of 2,846 mouse tumor samples derived from 212 diverse studies deposited in the NCBI’s Sequence Read Archive. We benchmarked these data against a previously established huTMEs classification^3^ built on TCGA^41^, the largest transcriptomic repository of bulk human tumor samples. Across this large mouse cohort, a majority of the samples (>60%) were classified as “Lymphocyte depleted” TME subtype (C4), defined by low numbers of leukocytes and a high macrophages-to-lymphocytes ratio^3^ (**Figure 2C**). Conversely, we observed a scarcity of mouse tumors classified within the TME subtypes characterized by high immune content (C2 and C6, comprising only ∼8% of the 2,846 mouse samples). To this extent, at least the bias towards lower infiltration seems to be a predominant trait of mice tumors, across most models.

We also sought to test whether the strong biases toward low T cells and high myeloid content could be due to an overrepresentation of heterotopic, cell-line based models in our particular selection of our murine cohort. We thus analyzed the T cell:Myeloid ratio across species and conditions (**Figure S1F-G**). This revealed that all huTMEs are biased toward T cells for composition while all muTMEs, including the spontaneous, slow-growing tumor lesions from MMTV-PyMT mice, are biased toward myeloid cells (**Figure S1F**). In addition, this ratio remained biased toward myeloid cells in KPC pancreatic tumors implanted either orthotopically or subcutaneously, as well as in B16, LLC and MC38 tumors inoculated to either mice housed under “dirty” conditions, which can promote T cell activation and lead to better modelling of human T cell responses^42^, and also to aged mice or to mice fed a high-fat ‘western’ diet (**Figure S1G**).

Taken together, these findings suggest that while muTMEs display model-driven diversity in their tumor-immune profiles, they overall represent the composition of only a modest fraction of huTMEs, primarily those resembling immune-desert, macrophage-rich TME types, even when considering a spectrum of tumor types, implantation site and other typical variables used in most cancer studies.

### Divergence of murine tumors from chemokine networks present in human TMEs

Chemokine networks are significant contributors to immune cell densities in tissues, since cells producing chemokines attract others to their neighborhoods^43^. We performed a systematic analysis of chemokine and receptor transcript expression^7,44,43^ within key cell populations of the huTMEs and muTMEs, to assess which cell types dominantly produced them, noting again that the two species have ∼200M years of evolutionary distance. For this we analyzed bulk RNA sequencing data from sorted T cells, Treg, myeloid, tumor and stroma compartments from huTMEs^7^, which we compared to pseudo-bulked scRNAseq datasets taken from our muTMEs cohort (**Figure S2A**). The higher resolution of the mouse scRNAseq also allowed us to subsequently assess cluster-specific expression of each chemokine (**Figures S2B** to **E** and **S3B** and **D**).

Given the prominent sparsity of T cells in muTMEs, we first analyzed the expression patterns of transcripts for the most prevalent chemokine receptors across each major compartment and species (**Figure S3A** and **C**), before focusing our analysis on prototypical T cell-specific chemokine receptors. This demonstrated significant variation—for example while transcripts encoding CCR2 and CCR5 are similarly expressed in the non-granulocytic myeloid compartment in both TMEs, they are more lowly expressed in T cells from the muTMEs compared to huTMEs (**Figure 3A-B**). This decrease was consistent in mouse T cells across models and we were able to verify the significant reduction in T cell expression of CCR5 at the protein level (**Figure 3C**). The relative reduction in CCR2 and CCR5 transcripts in aggregated T cells was also found when CD4 and CD8 T cells were analyzed separately (**Figure S3B**), suggesting that the reduction seen in muTMEs is not a result of a bias in their CD4:CD8 ratio compared to huTMEs. While these two chemokine receptors were notably skewed in their distribution, six other predominant T cell-biased chemokines receptors in the huTMEs were similarly highly T cell-biased in muTMEs (CCR4, CCR7, CXCR3, CXCR4, CXCR5, and CXCR6, **Figure 3A**), suggesting that this effect is not an artefact nor a global trend in all chemokines.

**Figure 3:**
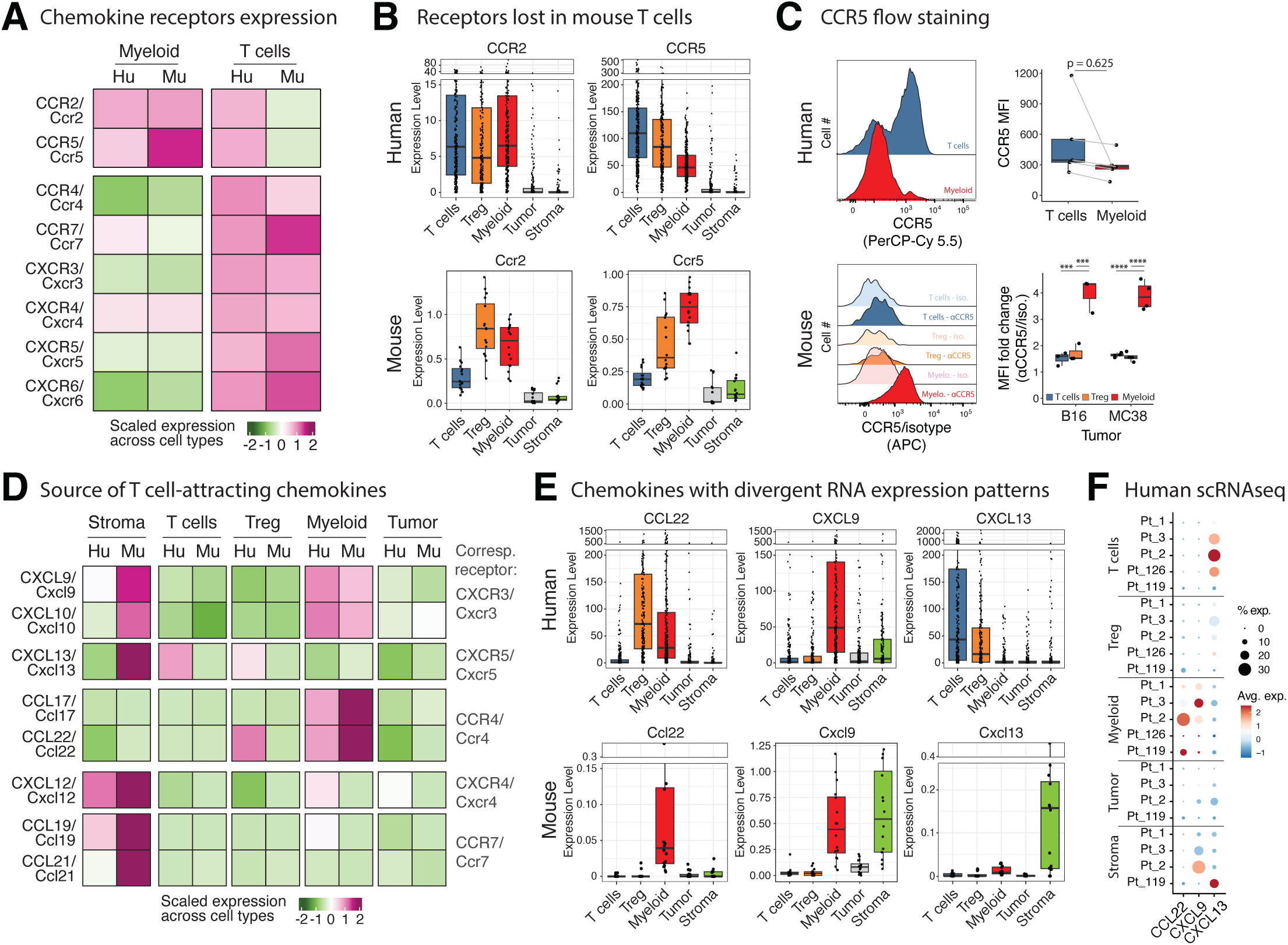
Divergent chemokine networks underly dysregulated T cell abundances in the tumor microenvironment. **A.** Heatmap comparing the scaled expression of chemokines receptors between human and murine T cells. For each species, the expression of each receptor was extracted for all T cells (bulk-sorted in human vs pseudo-bulked from scRNAseq in mouse) and scaled compared to their expression in the other available TME compartments of the same species (i.e Treg, myeloid, tumor and stroma). We then extracted the scaled values for T cells and plotted them side-by-side in a heatmap. **B.** Boxplots showing the expression levels of CCR2 and CCR5 across T cells, Treg, myeloid, tumor and stroma in human (top, shown as TPM from bulk RNAseq, each dot represents a single patient) vs murine tumors (bottom, shown as the expression level from scRNAseq averaged per sample, each dot representing a single mouse sample), exemplifying the differences found in A. **C.** Flow cytometry validation of the protein expression of CCR5 in human (top) and murine (bottom) tumors across different cell compartments, plotted as representative histograms (left) and boxplots of MFI (right). **D.** Heatmap (as in A.) comparing the expression of specific chemokines ligands binding the receptors found conserved in A. between the different cellular compartment of human vs murine tumors. **E.** Boxplots as in B. showing the expression levels of CCL22, CXCL9 and CXCL13 across cellular compartment and species. **F.** Dotplot presenting the scaled expression of CCL22, CXCL9 and CXCL13 across the cellular compartment of 3 different patients bearing HNSC tumors analyzed by scRNAseq. Statistical significance in C. was calculated using a t-test with Bonferroni correction, * p.adj ≤ 0.05, ** p.adj ≤ 0.01, *** p.adj ≤ 0.001, **** p.adj ≤ 0.0001.

We similarly analyzed inter-species variations in the cellular sources of the ligands for these six conserved receptors (**Figure S3C-D** and **Figure 3D**). Many T cell attracting chemokines were similarly distributed between huTMEs and muTMEs, notably those for CCR4, CCR7, CXCR4 and CXCR6 networks. For example, in both species CCL17/CCL22 and CXCL16 are consistently found expressed by myeloid cells, while CCL19/CCL21 and CXCL12 are predominantly expressed in non-immune stromal populations, except for high levels of CCL22 found in Treg from huTMEs but not muTMEs.

By contrast, chemokine networks involving CXCR3 and CXCR5, two major drivers of T cell infiltration in the TME^44,43,45,46^, were highly dissimilar. In muTMEs, CXCL9/CXCL10 and CXCL13 transcripts were significantly enriched in the stroma (i.e fibroblasts), while heavily biased towards myeloid and T cells/Treg respectively in huTMEs (**Figure 3E** and **Figure S3C-D**). This bias was not absolute since, by analyzing 5 individual human samples by scRNAseq, we observed that 1 in 5 samples displayed the muTMEs propensity for higher expression of CXCL13 in the stroma as compared to T cells (**Figure 3F**). This suggests that such a bias may not represent an inability of fibroblasts to produce CXCL13 in human, but more likely represents upstream signals that do not typically conspire to generate it in human fibroblasts populations. These particular observations regarding CXCL13 bear an immediate importance for the field, as recent studies have suggested that immune checkpoint responsive networks are organized around CXCL13+ T cells in patients^47–51^ and we find that rarely occurring in mouse models. We note the possibility that translation and secretion of these chemokines may be compensatorily regulated to achieve equivalent proteins levels from each transcript—a notoriously difficult thing to measure in intact TMEs. Nonetheless, these observations demonstrate an important fence and safety corridor for guiding future mouse studies of human-relevant transcriptomic networks, particularly those related to the assembly of ‘hubs’^5^ and prototypical archetypal distributions^7^.

### Interspecies deviations in TMEs immune cellular networks

Multiple studies have demonstrated that the presence of one cell type in the TME can correlate with another, typically because one either recruits or supports the other^31,52,53^. We therefore analyzed specific relationships in cell densities between cell types in huTMEs vs muTMEs, first examining correlations between population proportions and then examining these relationships across samples (**Figure 4**). Total immune infiltration inversely correlated with the frequency of tumor cell expressing the protein Ki67 (which allows passage through the G2 check point in the cell cycle, **Figure S4A**) in mice as well as human TMEs (**Figure 4A-B**). Similarly, both species exhibit positive correlations between CD8 T cells frequencies and the degree of exhaustion of those T cells (**Figure 4C**, **S4B**), as well as a relationship between cDC1/cDC2 frequencies versus CD8/CD4 T cell frequencies, indicative of co-maturation of these cells in tumors as previously described^30,54^ (**Figure S4C**).

**Figure 4:**
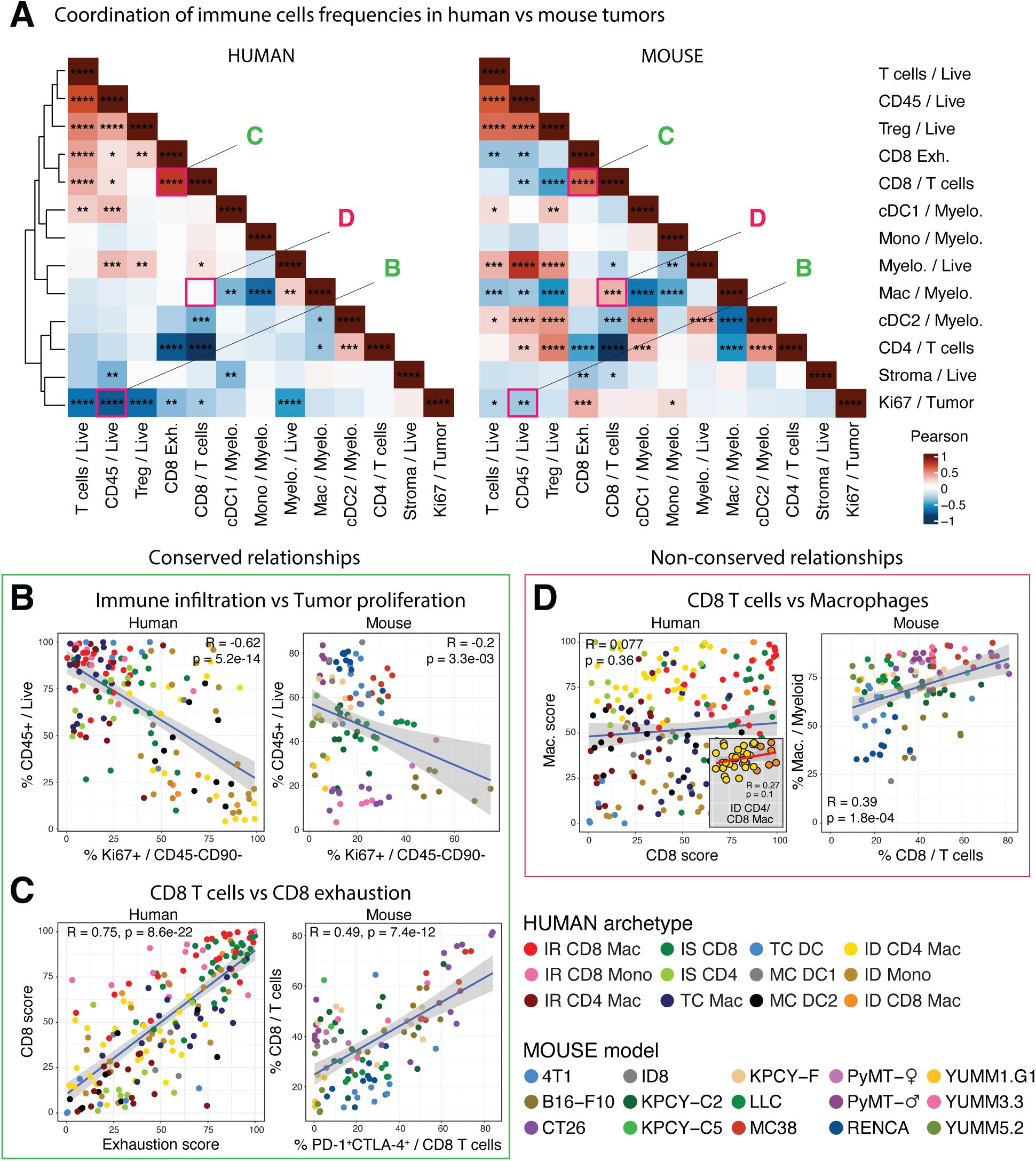
Differential conservation of relative cell densities in the tumor microenvironment. **A.** Pearson correlation matrices (human on the left, hierarchically clustered; mouse on the right, ordered according to the human matrix), presenting the correlations between the frequencies of different immune components in the tumor microenvironment. **B. to D.** Side-by-side dot plots exemplifying the correlations shown in A. of immune parameters between human (left, colored by archetypes) vs murine (right, colored by tumor line) tumors. The specific correlations are indicated above each panel and grouped as conserved correlation on the left (B. and C.) and non-conserved correlation on the right (D.). Statistical significance was calculated using a Pearson correlation test with Benjamini-Hochberg correction, * p.adj ≤ 0.05, ** p.adj ≤ 0.01, *** p.adj ≤ 0.001, **** p.adj ≤ 0.0001.

HuTMEs and muTMEs diverged with respect to a few well-described correlations between cell types. For example, CD4 Tconv frequencies correlated with regulatory CD4 Treg frequencies in muTMEs, but these were not well correlated in huTMEs (**Figure S4D**). More importantly, general correlations between T cells and non-granulocytic myeloid cells (**Figure S4E**) or more specifically between CD8 T cells and macrophages (**Figure 4D**) in muTMEs^31^ was not observed overall in huTMEs. However, when huTMEs were censored to only include the archetypes most similar to mice (immune deserts enriched in CD4/CD8 T cells and macrophages; **Figure 2A-B**), the correlations were restored (insets in **Figure 4D** and **S4D**). This may indicate that these cells track one another but only under specific conditions—namely in the absence of a large T cell pool and/or a global bias of the TME toward myeloid cells. These observations thus provide additional strong guardrails for interpreting murine tumor efficacy data for drug treatments which target these cell populations, and which anticipate conjugate interactions with these partner populations.

### cNMF identifies robust, cross-species transcriptomic programs

To examine the divergence and conservation of huTMEs vs muTMEs at a more granular level, and without pre-selection of features, we sought to use semi-supervised machine learning methods to discover fundamental gene programs occurring in each species, as well as co-occurring between cell types^24^. We thus processed both muTMEs and huTMEs transcriptomic data through a consensus non-negative matrix factorization analytic pipeline (cNMF^5,55–57)^, anchoring on T cells and non-granulocytic myeloid cells (**Figure 5A**). Collections of genes whose expression summarized to single gene factors across each species and each cell compartment were considered as gene expression programs (GEPs). These included GEPs that were overlapping between huTMEs and muTMEs in a given cell compartment, called ‘similar’ GEPs (**Figure 5**). Pairs of GEPs whose enrichment in one cell compartment was correlated with the enrichment of another GEP in another cell compartment were termed intercellular GEPs ‘movements’ (**Figure 6**).

**Figure 5:**
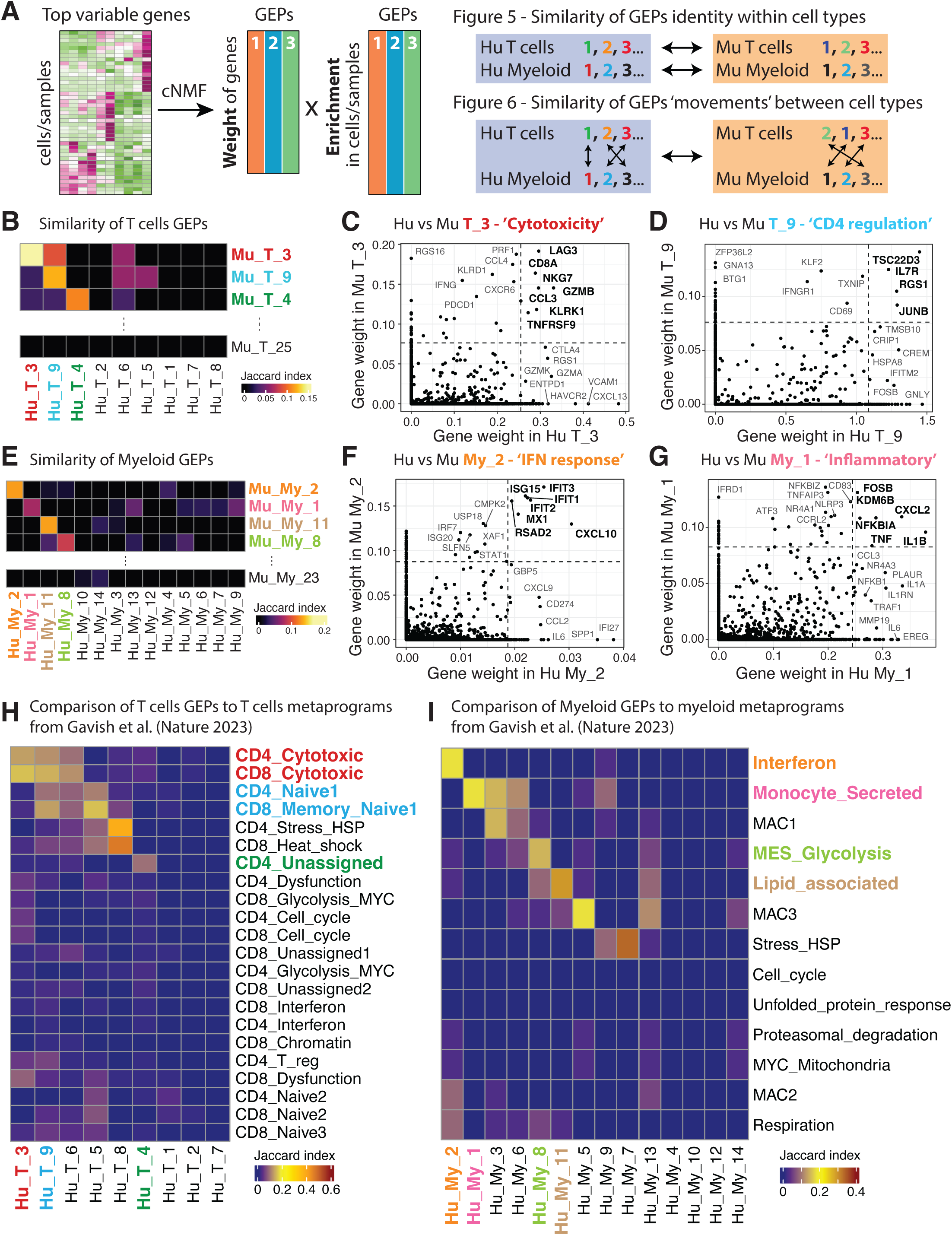
Isolation of conserved and robust gene expression programs in T cells and myeloid cells. **A.** Schematic of our analytical strategy using cNMF to compare the identity and coordination of gene expression programs (GEPs) in T cells and non-granulocytic myeloid cells across species. **B. and E.** Heatmaps showing the Jaccard indexes used to quantify the similarity between human vs mouse GEPs in T cells (B., using top 20 genes per GEP) and myeloid cells (E., using top 50 genes per GEP). Similarities of interest are highlighted in bold, colored fonts. **C., D., F. and G.** Scatter plots showing the genes contribution (i.e. genes weight) to human vs murine GEPs T_3 (C.), T_9 (D.), My_2 (F.) and My_1 (G.). Genes in bold show the highest overlapping contributions across species, while the genes in grey have a higher contribution in one species vs the other. The dashed lines separate the 40 highest contributor genes from the others in either factor. **H. and I.** Heatmaps showing Jaccard indexes used to quantify the similarity between our human T cells (left) and myeloid cells (right) GEPs to the ones published in^58^.

**Figure 6:**
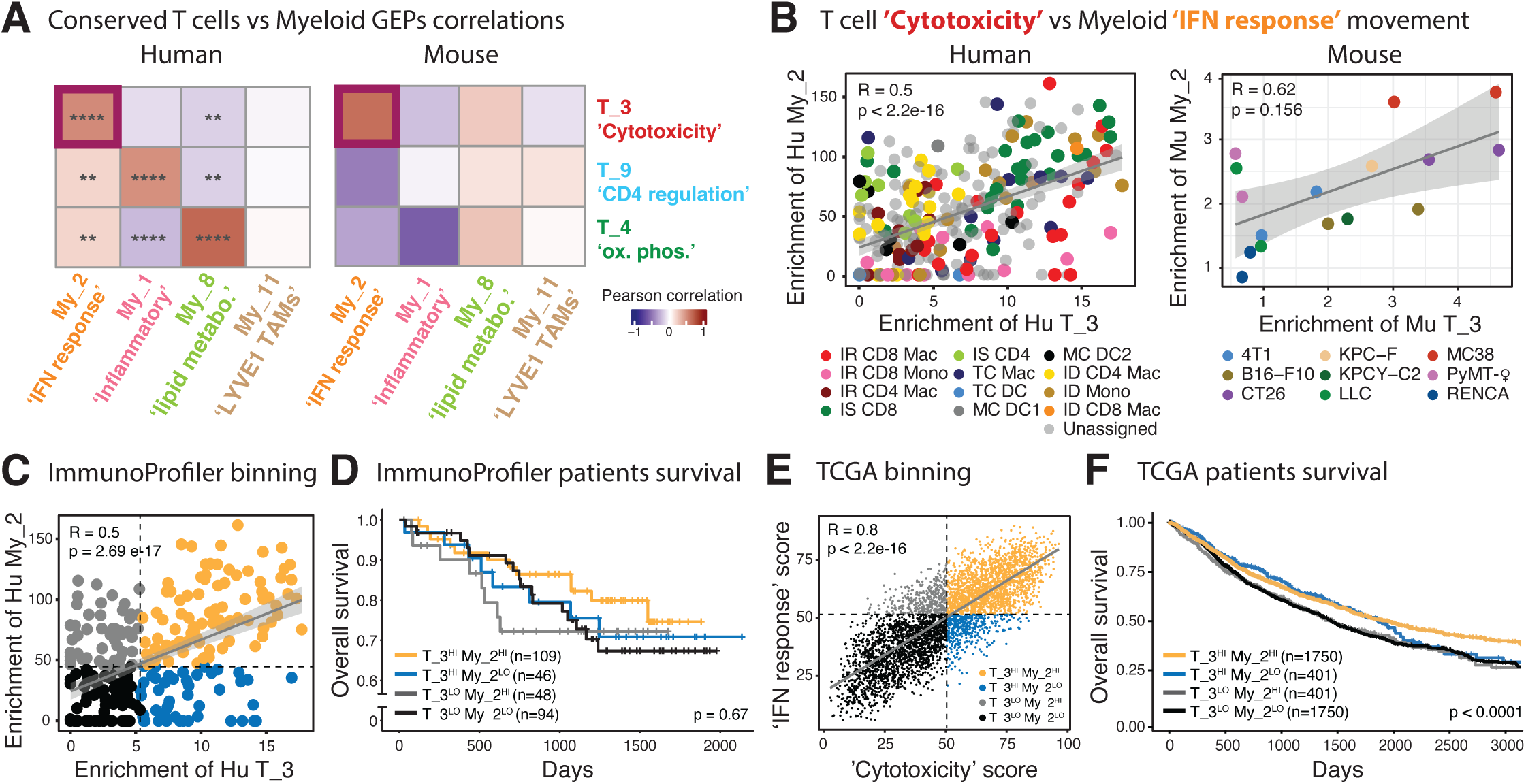
Cross-species conserved GEPs ‘movements’ between T cells and myeloid cells parses patients survival. **A.** Heatmaps presenting the Pearson correlations of T cells vs myeloid GEPs enrichment across human (left.) or murine (right.) TMEs. **B.** Scatter plots showing the correlation between the enrichments of GEPs T_3 and My_2 found across human (left, colored by archetypes) and mouse tumors (right, colored by tumors lines). Each dot represents a sample, and the diagonal grey lines represent linear regressions. **C.** Scatter plot (corresponding to left of B.) showing the binning of ImmunoProfiler patients as High or Low for T_3 and My_2 GEPs (respectively if present in the top or bottom 50% for each GEP enrichment). **D.** Kaplan-Meier graphs showing the overall survival of ImmunoProfiler patients stratified according to the binning shown in C. **E.** Scatter plot showing the relative enrichments of gene signatures calculated using the top 20 genes of GEPs T_3 and My_2 in TCGA patients. Patients were binned as High or Low for each GEP if they were present respectively in the top or bottom 50% for each calculated GEP gene score. Each dot represents a sample, and the diagonal grey line represents linear regression. **F.** Kaplan-Meier graph showing the overall survival of TCGA patients stratified according to the binning shown in E. Statistical significance in A., B., C. and E. was calculated using a Pearson correlation test with Benjamini-Hochberg correction. Statistical significance in D. and F. was calculated using a log-rank test.

When analyzing the T cell compartment in huTMEs (Tregs excluded^7^), we found nine stable T cell huGEPs (**Figure S5A** and **Table S1**). For purposes of naming these, we integrated information from existing gene ontology (GO) with known functions. Thus, huGEP T_3 containing genes such as PRF1, LAG3, GZMB, CCL3, CCL4 and NKG7 is tightly linked to ‘T cell cytotoxicity’, while T_9 is linked with ‘CD4 regulation’ (TSC22D3, JUNB, RGS1, IL7R, CD69) and T_4 to ‘oxidative phosphorylation’ (**Figure S5C**). The enrichment of some of these huGEPs was correlated with TME composition (**Figure S6A**). For example, ‘T cell cytotoxicity’ correlated with the frequencies of CD8 T cells, whereas T_5 ‘CD4 T cells-associated’ correlated profoundly with CD4 T cells frequencies. ‘CD4 regulation’ was correlated with stroma and Treg enrichment in the tumor, possibly representing a mechanism by which effector CD4 T cell regulation is driven towards inhibitory circuits in TMEs. Repeating this analysis in muTMEs uncovered twenty-five stable T cell GEPs (**Figure S5E**), similarly associated with T cells function (**Figure S5G**) and TME composition (**Figure S6D**).

To quantify cross-species overlap of these GEPs, we used a Jaccard similarity index and found high degrees of similarity in three out of the nine human T cell GEPs (**Figure 5B**, **S6G** and **I,** and **Table S1**). This notably included T_3 ‘T cell cytotoxicity’ (**Figure 5C**) and T_9 ‘CD4 regulation’ (**Figure 5D**), both similarly correlated with composition (**Figure S6A** and **D**). We speculate that when no homolog of a GEP was found across species (six out of nine huGEPs), this might either represent a limitation of the computational discovery method (starting from bulk RNAseq in human vs single-cell RNAseq in mouse) or an absence of that program as a major contributor.

Applying the same approach to the non-granulocytic myeloid compartment in huTMEs identified fourteen stable GEPs (**Figure S5B**). These included My_1 ‘inflammatory’ (IL1A, IL1B and NLRP3) and My_2 ‘IFN response’ (IFIT2, IFIT3, ISG15 and CXCL10) (**Figure S5D** and **Table S1**). Some of these again were highly correlated with cellular composition, including association of (i) conventional type I DC (cDC1) densities with My_10 ‘regulation of cytokines production’, (ii) tumor associated macrophages (TAMs) densities with both My_5 (‘chemotaxis’) and My_8 (‘lipid metabolism’) and (iii) monocytes densities with both My_6 (‘migration’) and My_9 (‘regulation of defense response’)(**Figure S6B)**. In muTMEs, we likewise identified twenty-three distinct and stable GEPs (**Figure S5F**) again often driven by genes known for their association with myeloid functions (**Figure S5H** to **J**) and/or whose enrichment correlated with TME composition (**Figure S6E**). Akin to T cells, a Jaccard analysis demonstrated high degrees of similarity between four out of the fourteen human myeloid GEPs in muTMEs (**Figure 5E** and **S6H**), including My_2 ‘IFN response’ (**Figure 5F**) and My_1 ‘inflammatory’ (**Figure 5G**) GEPs, as well as the TAMs-associated My_8 ‘lipid metabolism’ and My_11 ‘LYVE1 TAMs’ (**Figure S6J-K**).

To further assess the robustness of the conserved T cells and myeloid GEPs, we used a Jaccard analysis between these and an independent dataset describing various GEPs (or ‘metaprograms’) across populations, studies and tumor indications in human^58^. Doing so, we found that all our cross-species conserved huGEPs (e.g. My_1, My_2, My_8, My_11, T_3, T_4 and T_9) found an equivalent in this dataset (**Figure 5H-I**). This was also the case when comparing these GEPs to another recent study describing T cells and myeloid GEPs in glioma patients^59^ (**Figure S7A**), although sometimes our GEPs split into two in these other datasets, or we split a previously defined GEP into two in our analysis. We then asked whether these robust, conserved GEPs could also be observed in other biological processes than in tumors. We thus took advantage of a recent description of GEPs occurring over time and space in mouse myeloid cells during wound-healing^24^ and again applied Jaccard analysis approach (**Figure S7B**). This showed that three of these —My_1 ‘inflammatory’, My_2 ‘inflammatory’ and My_11 ‘LYVE1 TAMs’— had a similar GEP in the process of wound-healing (∼30% to 50% of genes overlapping for each similar GEPs between datasets).

This set of analyses overall defines a robust collection of core-conserved GEPs, alternatives and complements to ‘cell types’ or composition, which recur from mouse to human across studies and biological processes. We hypothesize that these may serve as modern measures of immune identity, in some cases providing metrics for describing overall immune status and similarity, with the cross-species conservation providing a tractable way to perturb and study the process in a model system.

### Coordinated movements of gene programs between immune cells in human vs mouse TMEs

We next assessed the degree to which, across species, the GEPs in one cell type were correlated with GEPs in another cell type, the concept of an intercellular GEP ‘movement’ or an indication of cellular crosstalk. For this we focused on the T cell-Myeloid axis as exemplar and because it is of known importance for antitumor immune responses^1,30,52,54^. We found a series of strongly correlated GEPs across cell types in huTMEs (**Figure 6A** and **S6C**), but that only one of these was also conserved in muTMEs (**Figure 6A** and **S6F**). This one comprised T_3 (‘T cell cytotoxicity’) and My_2 (‘IFN response’) GEPs, which were particularly correlated in the IR CD8 Mac, IS CD8 and ID CD8 Mac human archetypes and MC38, CT26 and B16-F10 mouse models (**Figure 6B**). To assess the relevance of this GEP ‘movement’ for patient outcome, we categorized patients as high/low for both T_3 (‘T cell cytotoxicity’) and My_2 (‘IFN response’) GEPs (**Figure 6C**) and observed how their combinations parsed out overall survival (**Figure 6D**). This showed a trend for improved survival of patients who displayed an overall high enrichment for both GEPs (T_3^Hi^My_2^Hi^) compared to any other combinations of these GEPs (**Figure 6D**).

Analyzing an independent transcriptomic dataset of whole huTMEs (TCGA^7,41^) demonstrated that these gene programs are also strongly correlated in this larger dataset (**Figure 6E**) —albeit extracted from entire tissue transcriptome. Utilizing the size of the TCGA allowed for a stronger parsing of the level of GEPs expression (high versus low) across a large number of patients with respect to outcome. In this analysis, high T_3 (‘T cell cytotoxicity’) generally associated with better outcomes as compared to lower, but T_3^Hi^My_2^Hi^ condition is the most favorable for outcome compared to all the other combinations (**Figure 6F**).

The constellation of genes in T_3—including granzymes but also IFNg itself—would thus appear to be broadly correlated with interferon-stimulated gene expression pattern (My_2) in myeloid populations, and that relationship appears to be highly conserved across mouse models. As described elsewhere^60,61^, we also found the relationship between cytotoxic T cells and interferon-stimulated myeloid cells to be beneficial for survival.

### Learning from discordant programs in human vs mouse TMEs

Multiple of the identified huTME intercellular GEP ‘movements’ that were found in huTMEs were not evidently conserved in muTMEs. For example, T_9 (‘CD4 regulation’) correlated positively with My_1 (‘inflammatory’) in huTMEs but these were poorly correlated in mouse (**Figure 6A** and **S7C-D**). We again noted that when we censored human samples to include only those most similar to muTMEs composition (ID CD4/CD8 Mac), that the correlation was less pronounced (**Figure S7C**), suggesting that the more mouse-like huTMEs had less of this correlated biology.

When this GEPs movement was analyzed in the TCGA dataset, we found the comprised gene programs were highly correlated (**Figure S7E**), and that patients with T_9^Hi^/My_1^Lo^ gene expression levels tend to have improved survival compared to other conditions (**Figure S7F**). CD4 T cell activation in the absence of concurrent IL-1-related myeloid inflammation—a typical concurrence in patients—could thus be optimal, consistent with previous reports associating poor outcome and inflammatory myeloid cells^62,63^. We also noted that the mouse-like archetypes (generally T_9^Lo^) were amongst the worst surviving, hinting that mouse models may be relevant systems to study these classes of poor survivors in patients.

Altogether, these analyses showcase the use of GEPs and their correlations as ‘movements’ as entry points to studying cellular crosstalk in muTMEs that is relevant to huTMEs and linked with patients outcome.

## Discussion

### Major findings

This study produced a data resource and systemic analyses examining the degree and possible circumstances to which muTMEs can represent and be studied to understand huTMEs. Our analyses revealed some definable specifics of the shortcomings that heretofore have not been molecularly defined. However, it also revealed swaths of similarities that we believe can be aligned with specific human tumors or can be studied as gene programs to understand principles related to therapeutic strategies.

Our data defines a number of metrics by which mouse models are apparently well-suited to study the immune response in macrophage-rich, poorly-infiltrated tumor archetypes (**Figures 1 and 2**) that span many tumor indications^7^. Some of the conserved relationships would appear intuitive, for example our finding that in the tumors of both human patients and mouse models a relationship exists between the degree of T cell infiltration and the degree of exhaustion of those infiltrating T cells. One may presume that such a relationship represents an axiom in which tumors cannot grow in the presence of high numbers of T cells if those are not simultaneously and proportionately desensitized.

Similarly, mouse models reinforce our previous finding^7^ that the level of Ki67 protein expression within human tumor cells is inversely correlated with T cell infiltration. Ki67 protein levels are highest during the G2 phase of the cell cycle and the protein is normally degraded upon successful mitosis^64^. Since huTMEs classified as ‘immune deserts’ and multiple mouse models have Ki67 fractions exceeding 25% of all tumor cells (**Figure 4**), we speculate that we may be observing a conserved tumor biology which fails to proteolyze Ki67 after mitosis, and which for as-yet-unknown reasons is not consistent with high immune infiltration.

Furthermore, some chemokines ligands-receptor networks involving CXCR4 and CXCR6 (**Figure 3**) are highly conserved from muTMEs to huTMEs. Other more complex themes exist across the spectrum of huTMEs and muTMEs, including coordination of large gene expression programs (GEPs, **Figure 5**), notably linking ‘T cell cytotoxicity’ and myeloid ‘IFN response’, that we and others have reported to be favorable to patients outcomes^65–67^.

Specifically in the context of such ‘desert’ huTMEs archetypes, murine models are also very relevant to study the coordination of specific immune cells at the population level, such as macrophages and their apparent close correlations with CD8 T cell densities (**Figure 4**), as well as some basic transcriptomic relationships between these cells.

In contrast, the majority of mouse tumors currently fall short at mimicking substantial biology of most huTMEs dominated by T cells, typically accounting for most of the bladder, kidney, lung or skin cancers^7^. This statement broadly applies to immune composition and coordination of population densities with one another, but also to the transcriptomic regulation of immune cells in these tumors. This last point is exemplified by the differential coordination of T_9 ‘CD4 regulation’ and My_1 ‘inflammatory’ myeloid cells between human and murine TMEs (**Figure 6** and **S7**) which warrants further examination as this myeloid GEP is generally linked with poor outcome^62,63^. As we study this conserved biology in greater detail—notably the value of IL-1 and/or TNF blockade—it may be that the patients that will benefit from cures derived in mice may be those with the worst current prognosis (**Figure 6**) since they similarly have the mouse-like combination of low overall immune abundance (**Figure 1**), high myeloid fractions (**Figure 2**), strong correlations between T cell and myeloid abundances (**Figure 4**) and poorly coordinated ‘CD4 regulation’/myeloid ‘inflammatory’ programs (**Figure 6** and **S7**).

While these conclusions fit 14 out of 15 of the mouse models we studied, the RENCA (‘renal carcinoma’) tumor model was an exception. This model is indeed highly infiltrated by immune cells, among which we find the highest amounts of myeloid cells, dendritic cells, as well as CD4 Tregs among our cohort of muTMEs (**Figure S1B**). RENCA seems to associate, although weakly, with human archetypes defined by a high myeloid content and biased toward CD4 T cells (IS CD4, MC DC1 and MC DC2^7^; **Figure 2B**). We also find RENCA to be the lowest mouse models for T_3 ‘T cell cytotoxicity’ and My_2 ‘IFN response’ GEPs enrichment (**Figure 6B**), while being the highest enriched for T_9 ‘CD4 regulation’ (**Figure S7D**). All these observations place RENCA as a model of choice to study immune populations and transcriptomic programs otherwise rarely found in other mouse tumors. Accordingly, our meta-analysis of 2846 public mouse samples also revealed that about ∼9% of these samples are associated with the two most highly infiltrated immune subtypes in human (C2 and C6 in **Figure 2C**). This provides a first example of a murine tumor that can better model a class of huTMEs and broadly confirms that mouse models can, in currently rare occasions, mimic a high degree of immune infiltration found in a large fraction of human tumors. Our resource thus provides a goal-line for the development and adoption of mouse tumor models that better mimic their human counterparts, in addition to providing insight into the ways in which mouse models may be better applied to study the very specific biology that they accurately replicate. Users may independently query this data at https://quipi.org/app/quipi_humu.

### Outstanding questions

Much work is still needed to provide a mechanistic explanation for the source of this general lack of TMEs similarity between human and mouse, defined by a paucity of T cells infiltration in murine tumors. It is possible that murine immune systems evolved to better respond to frequent infections and therefore mount better innate than adaptive responses (hence a paucity of T cells in mouse tumors). It is also possible that experimental models of inbred mice generally contain less T cells than wild mammals because of a lack of immune stimulation (pathogen infections, commensals colonization, frequent immunizations etc.) compared to human subjects. It has been shown that increasing microbial exposure to lab mice renders their immune systems, including T cell compartments, to better mimic those of humans^42,68,69^. Although we did not find this type of environmental normalization to affect significantly the T cell content of muTMEs (**Figure S1G**), this question deserves more careful exploration.

One potential source of TMEs divergence between mice and human might reside in the timescale of tumor growth. While it is thought that human malignancies often develop over many years, most experimental models in mice grow over a couple of weeks to several months. Even though there is no evidence to show that immune-rich/’hot’ tumors in human develop slower than immune-desertic/’cold’ ones, one could the hypothesize that the very fast growth of mouse tumors does not allow for a complex adaptive immune response to evolve. Embedded in this time component, we note that our study did not assess the dynamics of muTMEs composition. We purposedly analyzed our models at a transitional stage (day 14, most tumors measuring ∼300-500mm^3^) where tumors are consistent across model, large enough to be reliably measured and analyzed in depth but were not so large enough as to become overwhelmingly necrotic. Therefore, our study represents a snapshot of particular murine TMEs and did not capture the complexity of TMEs remodeling over time^20^, which could help reconcile some aspects of human vs murine tumor immunology.

At present, analysis of our data cannot explain the conserved or divergent mechanisms of response to immunotherapy between human and murine tumors. Given that the most anti-PD-1 responsive mouse models (for example, MC38) tend to be very poorly infiltrated by T cells and highly biased toward macrophages, and given the consideration that immunotherapy efficacy may involve both a de novo and pre-existing T cell response in human tumors^70–72^, it is likely that the benchmarking of responsive huTMEs against the responsive muTMEs will highlight even larger discrepancies than our study reveals. However, we think that a careful examination of transcriptomic coordination in the TMEs such as the one we introduce using cNMF-calculated GEPs ‘movements’ will be invaluable to establish the levels of conservation and divergence cross-species in response to specific therapeutic interventions. It is indeed likely that GEPs related to responsiveness or resistance to immunotherapy are already present in our dataset but will only be revealed as such when we will assess their relative enrichment after administration of therapy or various immune perturbations, ongoing work in our lab and now accessible using this data, for others with similar intent.

One major takeaway of our study given the recurring representation of specific human TMEs by mouse models at many levels, is that one may foresee that therapeutic strategies developed in murine systems (including their optimization using strategies mentioned above) will not be applicable to all patients. With understanding that the translation of therapies proven efficient in mice should be prioritized to historically treatment-resistant indications, there may be a silver lining. That is that those immune desert huTMEs biased toward high macrophages densities (for example gynecologic or pancreatic cancers), may be best represented by muTMEs in preclinical studies.

### Limitations

Our conclusions are based on the study of a non-exhaustive number of murine models, biased toward representing syngeneic tumor cell lines, and lacking a comprehensive analysis of all orthotopic, spontaneous, genetically engineered or xenograft tumor models. The few observations that we have made in such models (using MMTV-PyMT mice or KPC cell lines orthotopically) reinforced the general observations of a lack of global representation of huTMEs by mouse models regardless of their type of outgrowth or tissue of implantation (**Figure S1F-G**), but we anticipate that further analysis and studies will find other mouse tumors with features that better replicate the variety of huTMEs identified in comprehensive studies.

The degree of GEPs conservation that we find between human and murine immune cells may also be biased by the different sequencing platforms used to define these GEPs (bulk RNAseq in human versus single-cell RNA-seq in mouse). It is thus possible that more overlap exists between human and murine GEPs that we could not highlight in the present study. Nevertheless, we accounted for this by not over-concluding to the absence of similarity when no homolog of a GEP was found across species.

Relatedly, differences in the transcriptomic regulation (example of CXCL13 expression patterns) or coordination (example of T_9 vs My_1) that we observe between human and murine TMEs might arise from genetic divergence of promoter sequences and/or regulatory elements between human and murine genomes^11,73,74^. These differences may ultimately represent observed transcriptomic divergence of entire gene programs or result in a low degree of overlap between the top genes driving some human-murine similar GEPs (example of only 4 genes similarly driving T_9 in both species, highlighted in **Figure 5D**). This has notably been illustrated by the thousands of genes differently regulated by human versus murine mononuclear phagocytes when stimulated with lipopolysaccharide (LPS)^74^.

## Conclusions

Typical mouse models are imperfect representations of the spectrum of human tumors. While some cell-cell composition relationships and gene program ‘movements’ are well aligned mouse-human, notable ones are not, including cell-type biases in chemokine and chemokine receptor expression. The overall lesson of this study points to both the need and the potential for using more sophisticated analysis of how a drug pushes on the conserved circuits in a model, as a prelude to adopting muTMEs as surrogates for human cancers. In some cases (**Figure 4B-C**), cell-cell axes appear to be well-conserved. On the other hand, we discovered examples where censoring of certain human samples (**Figure 4D**, **S4E** and **S7C**) can select those huTMEs where muTMEs can model key relationships and biology. However, the chemokine networks that connect cellular composition within each species may be sufficiently diverged (**Figure 3**) and the resultant gene programs within subpopulations of cells sufficiently differentially biased (**Figure S6G-H**) to require caution when using mice to model therapeutic approaches for humans.

## Methods

### Human ImmunoProfiler samples

All human tumor samples were collected with patient consent after surgical resection under a UCSF IRB approved protocol (UCSF IRB# 20-31740), under the UCSF ImmunoProfiler project as described elsewhere^7^. Briefly, freshly digested tumor samples were analyzed by flow cytometry and/or FACS-sorted into conventional T cell, Treg, non-granulocytic myeloid, tumor and non-immune CD44+CD90+ stroma compartments to perform bulk RNA-seq on individual cell fractions.

### Mouse tumor models

#### Cell lines

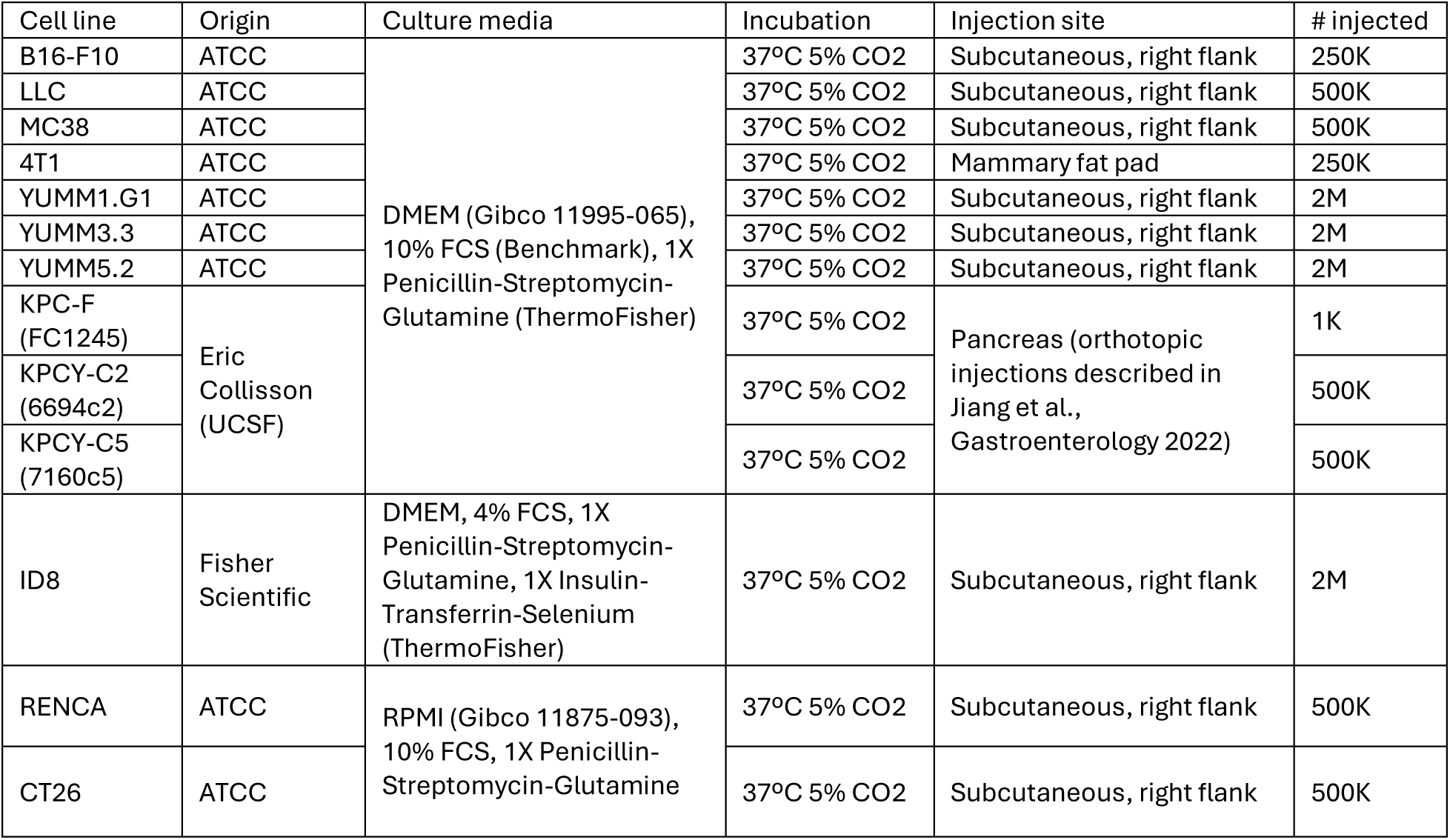

#### Animals

Unless specified, mice were housed at the AALAC-accredited animal facility of UCSF in specific pathogen-free (SPF) conditions, with typical light/dark cycles and standard chow. Animal experiments were approved and performed in accordance with the IACUC protocol number AN184232. For cell line-based models, 6-8 weeks old wild-type BALB/c (4T1, RENCA and CT26 models) or C57BL/6 (all other models) female mice were purchased from The Jackson Lab. Mice were euthanized at day 14 after tumor inoculation (typically ∼300-500mm^3^) for tumor harvest and further analyses. For high-fat diet experiments, mice were fed with either high-fat (Envigo TD.06414) or control (Envigo TD.08806) pellets for 2 weeks before starting tumor experiments and were kept on the diet during the 14 days of tumor course. For aging experiments, 90 weeks-old C57BL/6 female mice were purchased from The Jackson Lab and housed in our facility until reaching 2 years of age, when we started experiments. The genetically engineered mouse model (GEMM) MMTV-PyMT-mChOVA develop spontaneous tumor lesions from 4-6 months of age, specifically in the mammary gland in females and in the salivary glands in males^75^. In these models, mice were euthanized and tumors harvested when they reached ∼300-500mm^3^. In the experiments comparing mice housed under SPF or “dirty” conditions, as previously described^42^, age-matched 6-week-old C57BL6/J female mice were purchased from The Jackson Lab and split into two cohorts, one housed in SPF conditions and the other cohoused with pet-store mice, therefore considered ‘dirty’. Mice were housed/co-housed for 8-weeks prior to subcutaneous tumor cell injections of B16, MC38 or LLC, analysis of tumor growth, and endpoint analyses. Serologic dosage of antibodies specific to various pathogens was used to confirm the ‘dirty’ status of the co-housed mice.

### Sample preparation

After harvest, tumors were placed in a 12-well plate and minced to sub-millimeter pieces in 2mL of RPMI containing Collagenase IV (4 mg/mL) and DNase I (0.1mg/mL). Tissues were then incubated at 37°C for 30 min, with a mechanical dissociation step using thorough pipetting after the first 15 minutes. Digestion was stopped by adding 2mL of cold RPMI + 10% FCS + 1X PS-Glu to each sample before filtering through a 100 mm mesh and centrifugation at 500 g for 5 min at 4°C. Cells were then resuspended in RPMI + 10% FCS + 1X PS-Glu for further counting and analysis.

### Mass Cytometry

Mass cytometry (CyTOF) was performed as described elsewhere^76^. Briefly, conjugations of mass cytometry antibodies with metal isotopes were done using the Maxpar® conjugation kit (Fluidigm) according to manufacturer’s protocols and each antibody was titrated to define its optimal staining concentration. Each freshly digested sample was first stained with cisplatinium, fixed in 3.2% PFA and frozen at −80C. For CyTOF staining, the samples were then thawed and barcoded by mass-tag labelling with distinct combinations of stable Pd isotopes in 0.02% saponin in PBS before further pooling and staining. For this, cells were first resuspended in cell-staining media (Fluidigm) containing metal-labeled antibodies against CD16/32 for 5 min at room temperature to block Fc receptors, followed by the addition of a cocktail containing surface markers antibodies in a final volume of 500µL for 30 min at room temperature. Cells were then permeabilized with methanol for 10 min at 4 °C, washed and incubated with a cocktail containing intracellular markers antibodies in a final volume of 500µL for 30 min at room temperature (all antibodies listed in **Table S2**). Cells were finally stained with 191/193Ir DNA intercalator (Fluidigm) diluted in PBS with 1.6% PFA 48h prior to data acquisition. For acquisition, cells were washed and resuspended at 1M/mL in deionized water + 10% EQ four element calibration beads (Fluidigm) and analyzed on a CyTOF mass cytometer (Fluidigm). We acquired an average of 1–3 × 10^5^ cells per sample, consistent with generally accepted practices in the field. After data collection, we used the Premessa pipeline (https://github.com/ParkerICI/premessa) to normalize data and deconvolute individual samples. We then manually gated the individual FCS files using FlowJo (BD) according to the gating scheme described in **Figure S1A**.

### Comparison of human and mouse TMEs diversity

Mouse CyTOF data and human flow cytometry data (inferred using linearity of the flow parameters with gene scores as shown in^7^) were used to generate immune feature matrices based on the 10 PanCan immune features as previously described. After aggregating mouse samples by tumor line and human samples by immune archetype, the median frequency values for each immune feature were computed. Cosine similarity was computed by normalizing each row vector to unit length and taking the matrix product of the normalized data with its transpose. Cosine distance was then derived by subtracting the similarity values from 1.

### Immune subtypes classification of mouse tumors

All RNA-Seq samples used in this analysis were derived from mouse tumors and collected from the NCBI Sequence Read Archive (SRA). The following steps were followed to establish a cohort of mouse tumor samples. First, cancer-related disease ontologies (DOIDs) were gathered from the disease ontology database^77^ using the “ontologyIndex” package with search terms “cancer”, “carcinoma”, and “neoplasm”, resulting in 1,529 unique DOIDs. MetaSRA^78^, which maps SRA samples to terms in biomedical ontologies, was queried using the following parameters: “RNA-Seq”, “mouse”, “tissue”, and the cancer-related disease ontology DOIDs. Samples with single-cell RNA sequencing ontologies were excluded. This query to MetaSRA returned 3,900 samples, with incomplete metadata and SRA Sample IDs (SRSs). Next, MetaSRA results were filtered by manual examination of study metadata and abstracts to exclude samples not derived from tumors, which resulted in a final cohort of 2,846 samples from 212 studies used in the analysis. RNA-seq gene counts for each study were accessed through recount3^79^, ensuring the application of a uniform alignment pipeline and annotation records. Gene counts were normalized by dividing by the 75th percentile gene count for each sample and then applying a log2 transformation. Each mouse gene was converted to its human ortholog using the Babelgene package. To predict a tumor immune subtype for each sample we used ImmuneSubtypeClassifier^80^. This tool is a machine-learning model that compares quantile and gene-pair features of 485 genes to determine immune subtypes, as defined by^3^. Each sample was assigned its “BestCall” subtype.

### Mouse scRNAseq data generation

For most mouse experiment, we started by sampling 1e6 cells from the tumor of each animal and generated a single pool for each group. A group of 5 mice therefore generated a pool of 5e6 cells. This cell pool was then stained with the Zombie NIR viability dye (1/1000 in PBS, 10min at 4C), before being incubated with Fc block (clone 2.4G2, Tonbo Biosciences) and barcoded with HTO antibodies (TotalSeq-A from BioLegend). We then pooled all barcoded samples together and stained them with mix of fluorescent-labelled antibodies (**Table S2**). Using a BD FACSAria II cell sorter (BD Biosciences), we then gated live immune cells (Zombie-CD45+) and sorted 2 pools of cells from these samples: a pool of lymphoid cells containing equal amounts of T cells (CD90.2+), B cells (B220+MHCII+) and NK cells (CD49b+), and another pool of myeloid cells gated as CD11b+ and/or CD11c+ among the non-T-B-NK cells. These two pools were then washed, counted and then individually encapsulated following 10X Genomics specifications for v.3 3′ chemistry. Single cell cDNA library construction was then performed as directed by 10X Genomics standard procedures. Following fragment analysis and library quantification using the BioAnalyzer, the libraries were finally sequenced on Illumina NovaSeq SP using 10X Genomics recommended sequencing parameters.

### Mouse scRNAseq data processing

BCL files were converted to FASTQ format using cellranger mkfastq (v3.0.2). FASTQs were then aligned to the GRCm38 genome, generating gene-by-cell count matrices with cellranger count. The resulting matrices were imported into Seurat (v4.0.3) for downstream preprocessing and HTO demultiplexing, with manual inflection values provided to confidently retain only singlet cells (https://github.com/UCSF-DSCOLAB/aarao_scripts/). After demultiplexing, quality control scores were calculated for mitochondrial, ribosomal, and cell-cycling related expression. Low-quality cells were defined as those with >10% mitochondrial RNA content, >60% ribosomal RNA content, or fewer than 250 detected genes. Unless otherwise stated, the following processes took function defaults. Data were normalized (Seurat::NormalizeData), 3000 variable features selected (Seurat::FindVariableFeatures), and then scaled (Seurat::ScaleData) with regression against percent mitochondrial, percent ribosomal, and cell cycle S and G2M scores. Principle components 1:30 were calculated (Seraut::RunPCA). Data were merged across samples and batch corrected with (harmony::RunHarmony v0.1.1), using batch as the grouping variable and retaining all default parameters for harmonization. UMAP (Seurat::RunUMAP) was then calculated using 30 principle components with harmony selected as the reduction. Finally, cell type annotation was perfumed by identifying cell clusters (using Seurat::FindClusters) and grouping them based on their identify, inferred using their differentially expressed genes (using Seurat::FindAllMarkers with test.use=“poisson” and latent.vars = “Tumor_Line”) and prior knowledge of cellular markers. Note that during this process we identified clusters of non-immune cells, likely representing contamination during cell sorting, and containing both tumor and stromal cells. These were present at stable ratios across all experiments and we therefore decided to include the subpopulations representing PanCan-relevant tumor and stroma compartments^7^ for the chemokines analyses of **Figure 2**.

### Chemokines analyses

To process the human bulk RNA sequencing data, raw counts were filtered as previously described^7^. Filtered counts were normalized to account for differences in sequencing depth via TMM normalization, and converted to counts per million. Next, values were log transformed and then z-scored across all compartments, enabling comparative analysis of chemokine expression across different compartments and patient archetypes. To better compare our single cell RNA sequencing mouse data to the previously published bulk RNA sequencing data, the single-cell data were pseudo-bulked by calculating the average gene expression across each gene and condition and we then scaled the averaged expressions across all compartments. Finally, to identify homologous chemokines between mouse and human, the BioMart archive (April 2018) was used to convert mouse and human genes.

### Human scRNAseq

The cohort of samples used in **Figure 3** to analyze population-specific expression of human chemokines ligands in human HNSC tumors has already been described elsewhere^81^. Briefly, ImmunoProfiler tumor samples were digested to single cells and then submitted to single-cell gene expression analyses through the CITE-seq pipeline (Illumina) before analysis, coarse annotation of TMEs compartments and visualization of gene expression across patients and TMEs compartments.

### Data accessibility

Both human (https://quipi.org/app/quipi) and murine (https://quipi.org/app/quipi_humu) datasets can be readily queried and visualized using our user interface and reviewers are invited to visit those beta sites with the provision that they please not share them widely prior to publication of our study. Relevant raw data will be made publicly available at or before final publication.

### cNMF (consensus Non-negative Matrix Factorization)

The human dataset used in these analyses was a subset of the UCSF ImmunoProfiler Initiative (IPI) dataset described in^7^. We used bulk RNAseq gene TPMs (transcripts per million) from sorted conventional T cells and non-granulocytic myeloid cells. We filtered the data to include only samples from primary or metastatic tissue with a quality metric EHK score ≥ 8 (355 T cell samples and 322 myeloid samples). We applied log2 normalization to handle outliers and then filtered the genes to the 5000 most variable genes based on median absolute deviation (MAD) to reduce noise. To obtain a quantitative measure of NMF rank stability, we used the cophenetic correlation coefficient as described in^82^. We ran the NMF algorithm 50 times for each rank in the range of 3 to 30 factors. For each run, we generated a connectivity matrix, where the entries were 1 if the sample belonged to the same factor across runs and 0 if it did not. We then computed a consensus matrix by averaging all the connectivity matrices from the repeated runs. The cophenetic correlation coefficient, defined as the Pearson correlation between the consensus matrix and a sample distance matrix calculated using the Euclidean linkage method, was calculated for each rank. We repeated this process five times and plotted the median cophenetic correlation coefficient against the rank. Ranks that corresponded to local maxima in the median cophenetic correlation plot were identified as stable candidate ranks for further analysis. To obtain W and H matrices for the candidate ranks, we used a method similar to the single-cell workflow called DECIPHER-seq^57^. We ran the NMF algorithm 10 times for each of the selected ranks using the Python module sklearn^41^. We then concatenated the W and H matrices from all 10 runs to create larger matrices, W_10_ and H_10_. We then applied K-means clustering with k = candidate rank H_10_ matrix and used the resulting cluster labels on the W_10_ matrix. Next, we removed outliers in the clusters using the Local Outlier Factor (LOF) algorithm for anomaly detection, which computes the local density deviation of data points relative to their neighbors, applying a 40% cutoff. Removing outliers results in more homogeneous clusters and helps with the interpretation of results downstream. After outlier removal, we calculated the median W_10_ and H_10_ to form consensus matrices, W_Con_ and H_Con_.

For the murine dataset cNMF was ran using DECIPHER-seq^57^ on our mouse single-cell RNAseq data. Briefly, we isolated all conventional T and non-granulocytic myeloid cells and ran the DECIPHER-seq function ‘iNMF_ksweep()’ with parameter batch=FALSE to run NMF on each population for twenty replicates of each k from 2 to 40. Individual biological samples were used as the sample grouping variable.

### TCGA cohort and survival analyses

Analyses using the TCGA dataset was performed using the TCGA sub-cohort described in^7^. Briefly, tumor RNAseq counts and TPM along with curated clinical data for 13 cancer types (BLCA, COAD, GBM, GYN (grouping OV, UCEC and UCS), HNSC, KIRC, LIHC, LUAD, PAAD, SARC and SKCM) was filtered down to include primary solid tumors and metastatic samples only. This reduced the TCGA sample set to 4341 tumor samples. GEP scores were generated by first normalizing (using percentiles) the expression values of the top 20 contributor genes for each GEP across all patients, followed by averaging these 20 normalized values for each patient. For survival analysis, patients were categorized as either high (top 30%) vs low (bottom 30%) for each GEP score and analyzed using a log-rank test. The PanCan myeloid feature scores (relative to monocytes, macrophages, cDC1 and cDC2) were calculated in the same way, using the feature gene signatures from^7^.

## Acknowledgements

We thank Matt Spitzer (UCSF) and Peter Turnbaugh (UCSF) for thoughtful discussions about the project. We are grateful to Eric Colisson (UCSF) and Honglin Jiang (UCSF) for providing KPC tumors-bearing mice used in this study. We also thank Iliana Tenvooren (UCSF) and Stanley Tamaki (UCSF, Flow CoLab) for their precious assistance with CyTOF-related matters, Ravi Patel (UCSF, Data Science CoLab) for his input to data processing and Catherine Chu and Annie Poon (both from UCSF Genomics CoLab) for their help with scRNAseq quality control and libraries sequencing.

## Authors contribution

Conceptualization: TC, MFK

Experimentation: TC, NWC, GCR, JT, NVG

Data processing and analysis: TC, RGJ, BS, EF, AR, DB, LLJ, JPG, DAS, XV

Data visualization: TC, RGJ, BS, HW

Manuscript drafting: TC, MFK, RGJ, BS, Supervision: MFK, AJC, GKF

External collaborations supervision: DM, ETL

## Data availability

Both human (https://quipi.org/app/quipi) and murine (https://quipi.org/app/quipi_humu) datasets can be readily queried and visualized using our user interface and reviewers are invited to visit those beta sites with the provision that they please not share them widely prior to publication of our study.

## Supplementary data legends

**Table S1: Top 50 genes, ranked by weight, contributing to each human and murine gene expression programs (GEPs) in T cells (tab 1) and non-granulocytic myeloid cells (tab 2).**

**Table S2: Lists of antibodies used in our study.**

**Figure S1:**
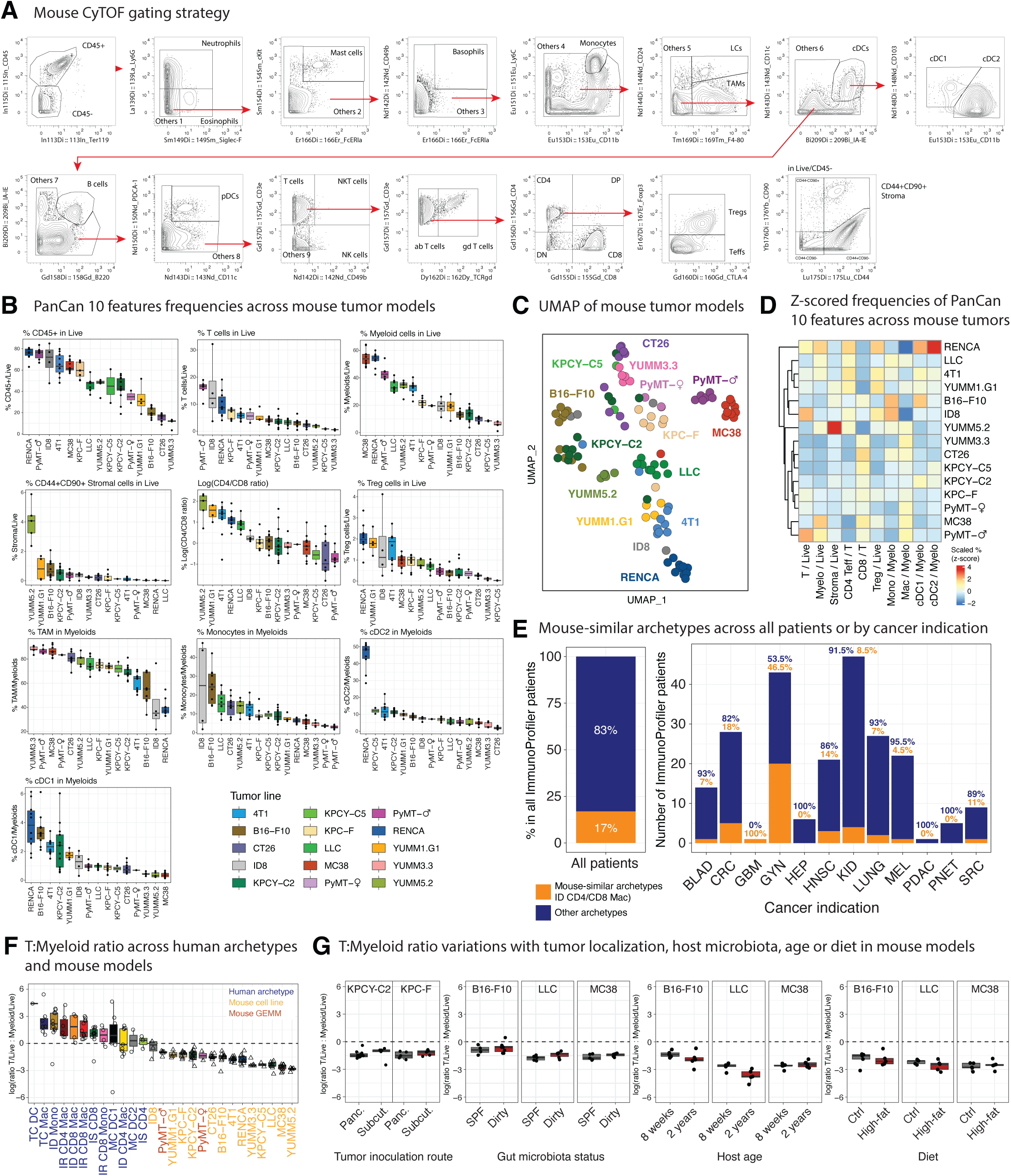
CyTOF gating strategy and diversity of immune profiles in muTMEs. **A.** Gating strategy of immune populations using CyTOF analyses of mouse tumors. **B.** Waterfall boxplots describing the specific frequencies of the PanCan 10 features across mouse models. **C.** UMAP dimensionality reduction of individual mouse tumor samples, colored by model, using the 10 PanCan features. **D.** Heatmap presenting the hierarchical clustering of mouse tumors using the scaled PanCan 10 features. Scaling consisted in calculating the z-scores of each feature across mouse samples before averaging per model and then plotting values. **E.** Bar graphs presenting the frequencies of human samples whose TME composition can be mimicked by mouse models (in orange, i.e belonging to ID CD4/CD8 Mac archetypes) or not (in blue), either across the whole cohort (left) or split by tumor indication (right). **F and G.** Boxplots presenting the ratio of T cells over myeloid cells frequencies in Live across human and murine tumors (F.), between orthotopically (Panc.) vs subcutaneously (Subcut.) injected KPC pancreatic tumor models (left of G.) or in B16, LLC and MC38 tumors grown in mice kept under specific pathogen-free (SPF) vs dirty housing conditions (center-left of G.), or grown in young (8-weeks) vs aged (2-years) mice (center-right of G.), or in mice fed with a control (Ctrl) vs High-fat diet (right of G.).

**Figure S2:**
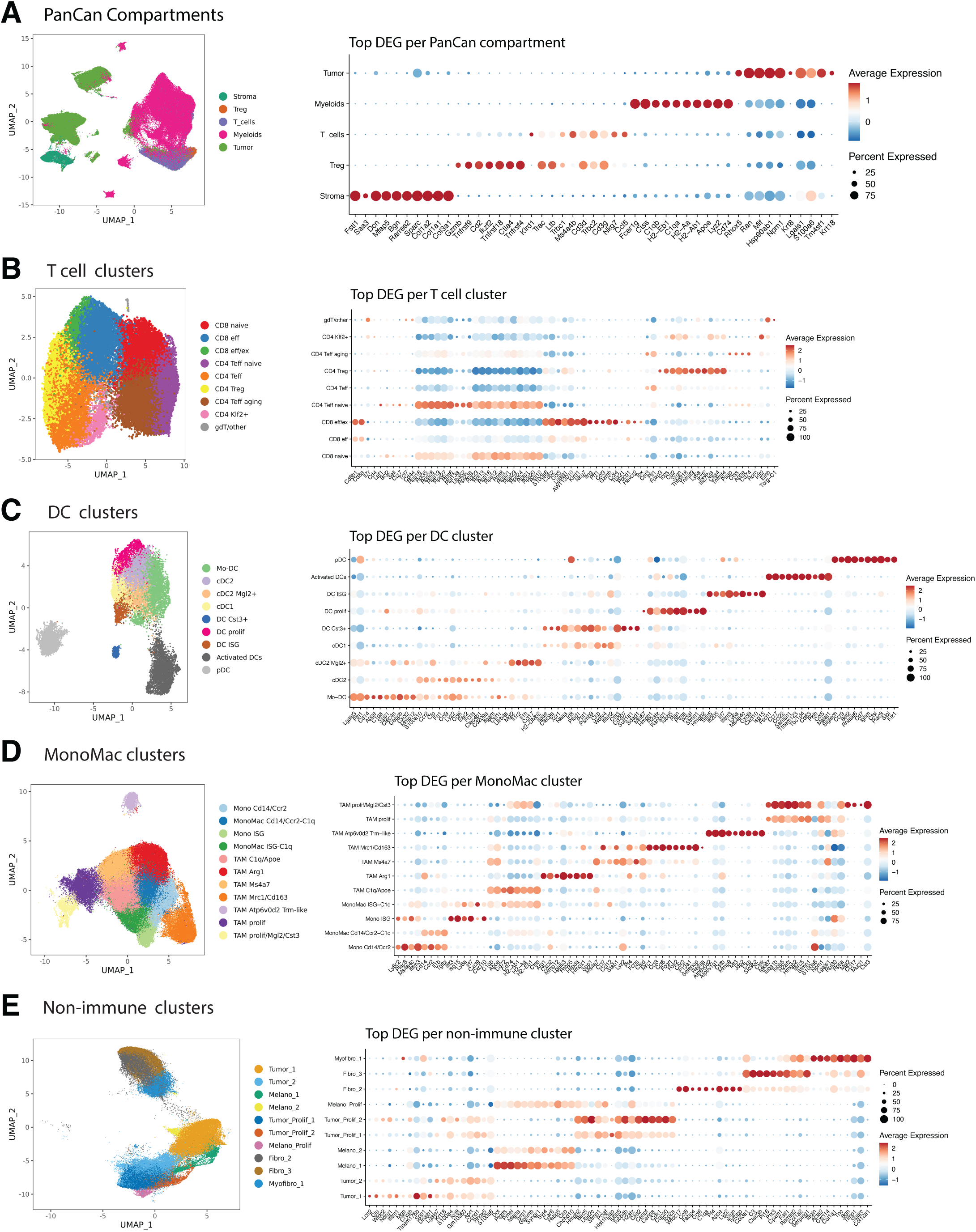
Murine scRNAseq dataset. **A.** UMAP visualization of 59551 cells merged from the entire mouse tumor scRNAseq data set (left), colored by their cellular compartment identity (i.e T cells, Treg, myeloid, tumor or stroma) and dotplot (right) presenting the scaled expression of the top 10 differentially expressed genes (DEGs) between the 5 compartments **B. to E.** As in A., UMAP plots (left) and dotplots of top DEGs (right) describing the specific clusters of murine T cells (B.), DCs (C.), monocytes/macrophages (i.e. MonoMac, D.) and non-immune cells (E.).

**Figure S3:**
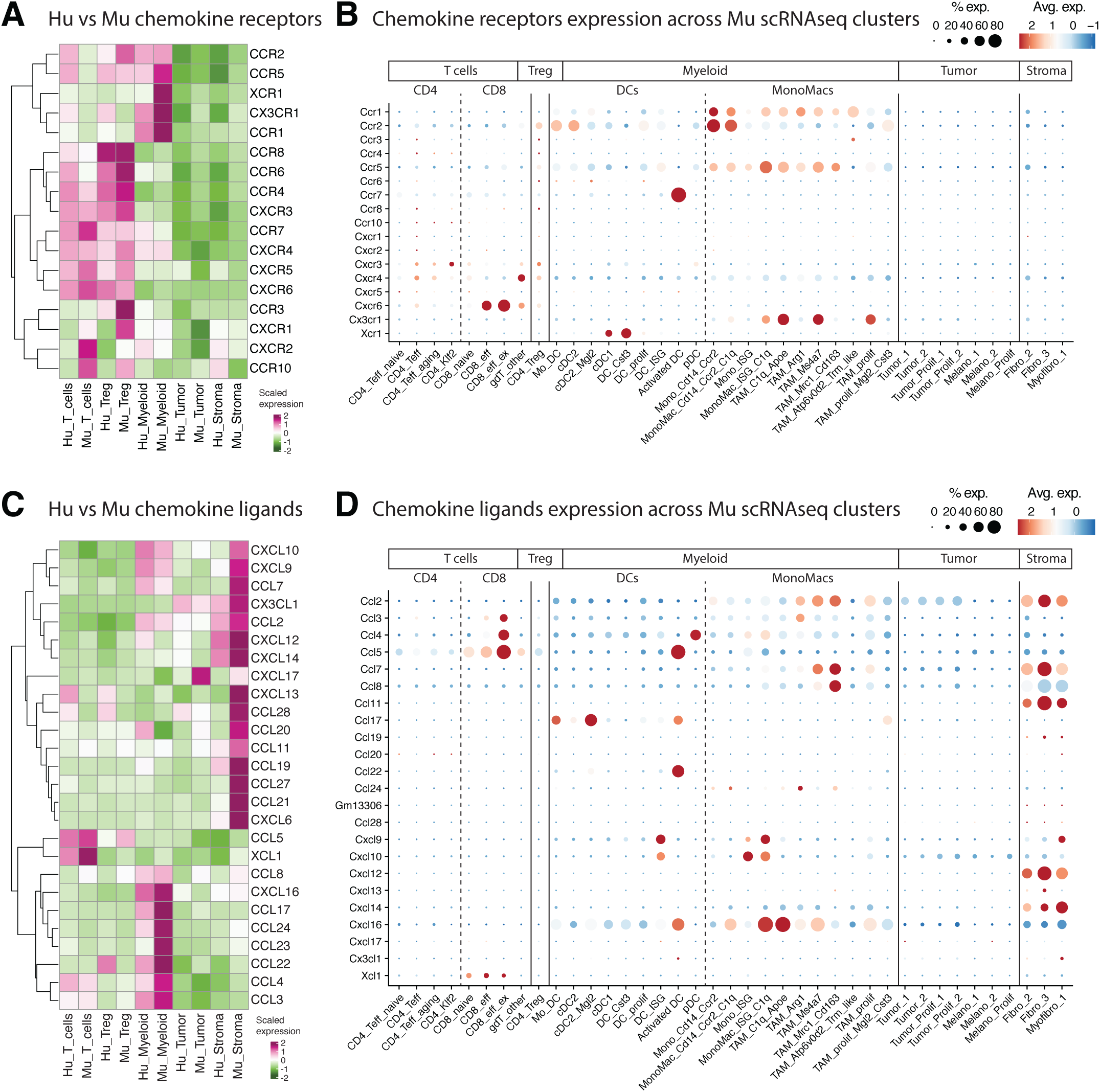
Additional chemokine pathways analyses. A. and. **C.** Heatmaps comparing the scaled expression of chemokines receptors (A.) and ligands (C.) between human and murine T cells, Treg, myeloid, tumor and stroma. For each species, the expression of each receptor/ligand was first scaled across compartments. We then extracted the scaled values for all human and murine compartments and plotted them side-by-side in a hierarchically clustered heatmap. **B. and D.** Dotplots presenting the scaled expression of the chemokine receptors (B.) and ligands (D.) shown in A. and C. across the murine cell clusters from scRNAseq shown in Figure S2B to E.

**Figure S4:**
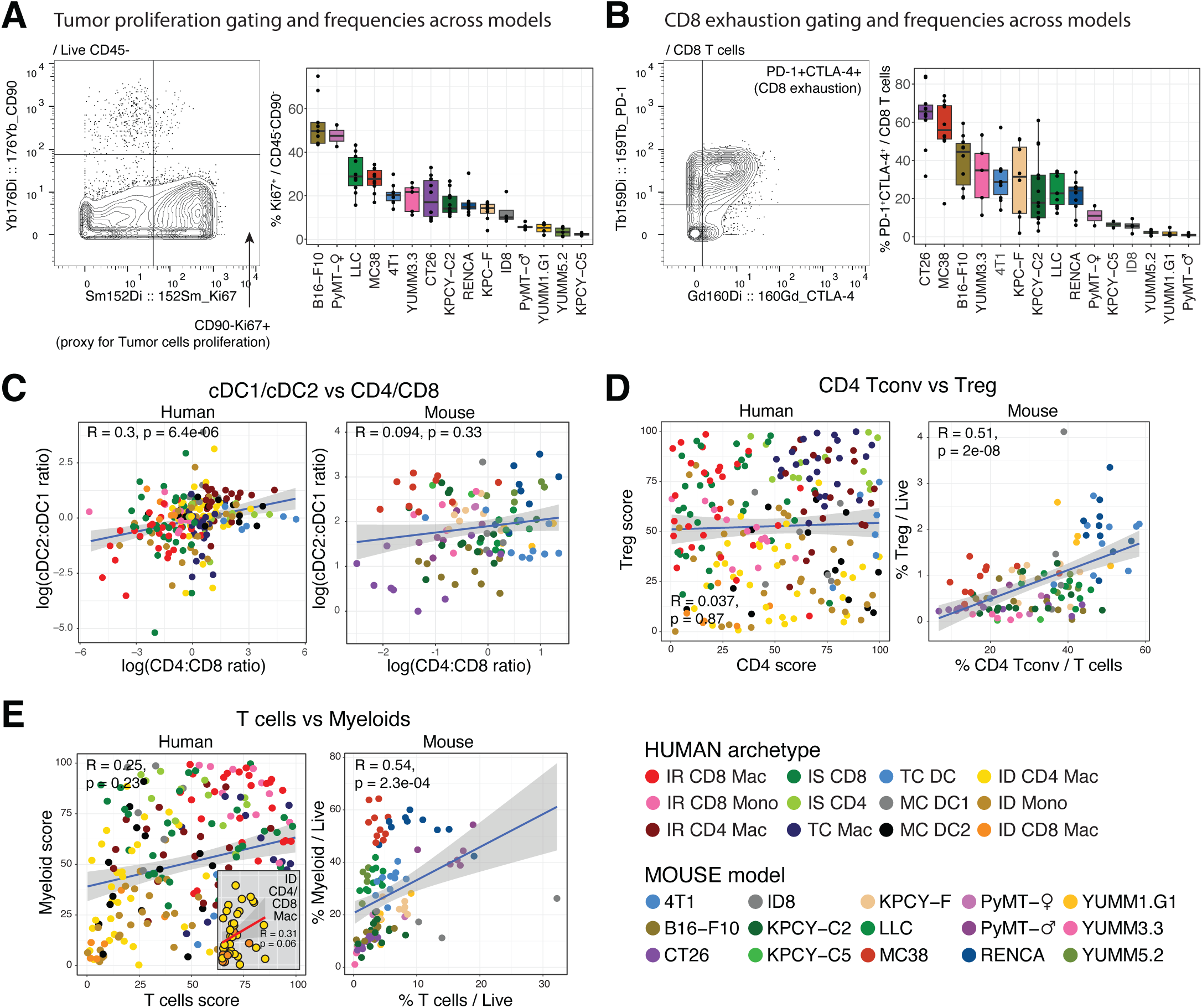
Additional examples of immune frequencies coordination. A. and. **B.** Gating strategy (left) and boxplot (right) presenting the frequencies of tumor proliferation (defined as Ki67^+^ in CD45^-^ CD90^-^ cells, A.) and CD8 exhaustion (defined as PD-1^+^CTLA-4^+^ in CD8 T cells, B.) in mouse tumor samples. **C. to E.** Dot plots exemplifying some correlations shown in Figure 4A. between human (left, colored by archetypes) vs murine (right, colored by tumor line) tumors. Statistical significance in C. to E. was calculated using a Pearson correlation test with Benjamini-Hochberg correction.

**Figure S5:**
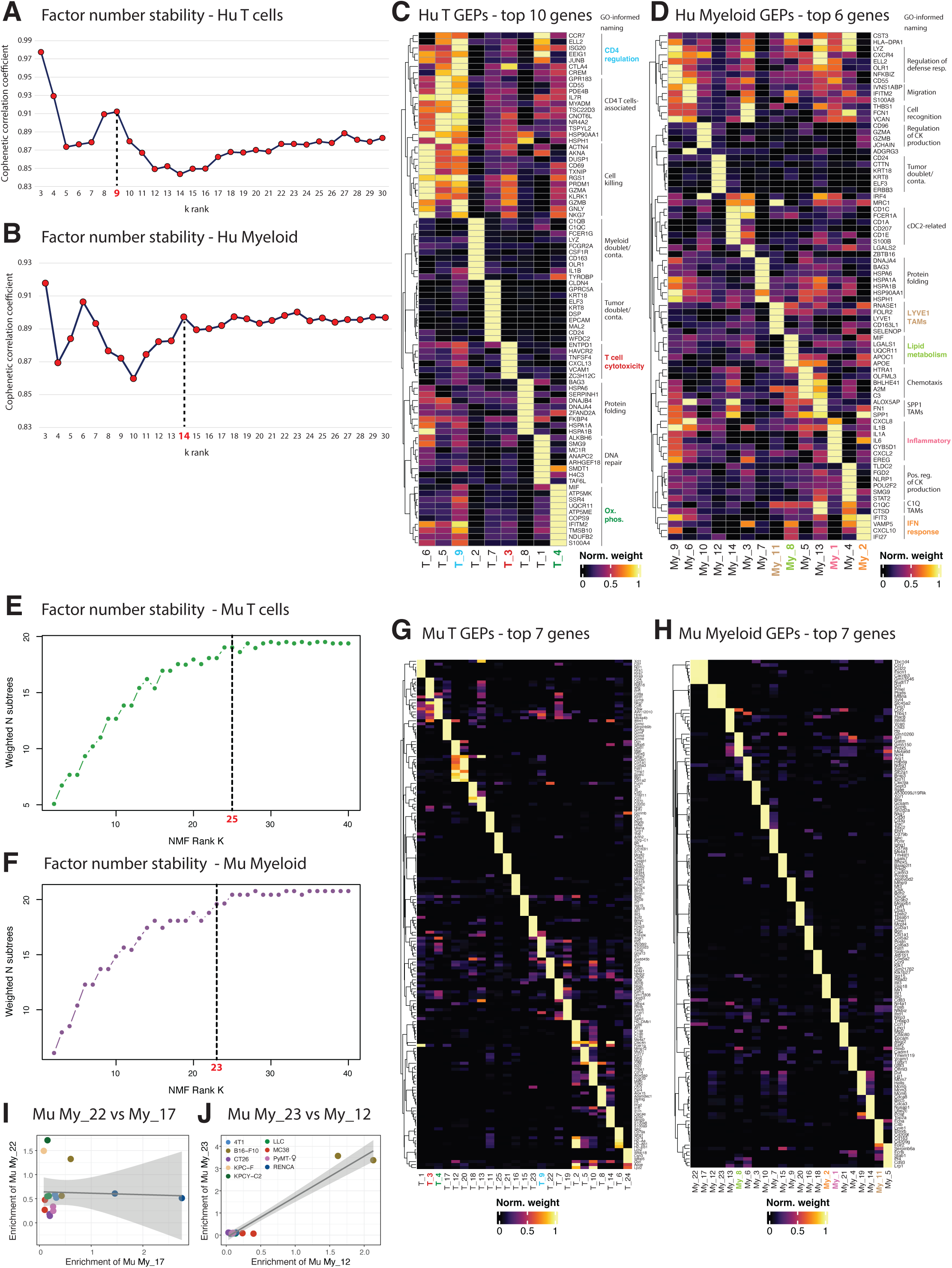
Identification of T cells/myeloid gene expression programs (GEPs) using cNMF. A. and. **B.** Scatter plots showing the cophenetic correlation coefficients (CCC) used to establish the stability of each cNMF resolution in human T cells (A.) and myeloid cells (B.). We picked the number of stable GEPs (in red) as the value associated to a peak in CCC followed by a flattening of the curve. **C. and D.** Heatmaps showing the scaled weights (scaling first all genes in each GEP and then each scaled gene across all GEPs) of the top genes composing T cells (C.) or myeloid (D.) GEPs. **E. and F.** Scatter plots showing the weighted N subtrees (WNS) values used to establish the stability of each cNMF resolution in mouse T cells (E.) and myeloid (F.). We picked the number of stable GEPs (in red) as the value associated to the highest WNS value that was followed by a flattening of the curve. **G. and H.** Heatmaps as in C. and D. showing the scaled weights of the top genes composing mouse T cells (G.) and myeloid (H.) GEPs. **I. and J.** Scatter plots showing the relative enrichments of GEPs My_17 vs My_22 (I.) and My_12 vs My_23 (J.) across mouse samples, colored by tumor model. Note that among the 23 murine myeloid GEPs we noticed 2 apparent factors duplication, namely My_22/My_17 and My_23/My_12. While My_22 and My_17 seem to share gene composition (related to activated DCs), they are enriched in different sets of samples (I.). However, My_23 and My_12 appear as a duplication that could arise from over clustering of the NMF algorithm, because their gene composition shows a low degree of contamination/doublet with melanocytes and they are both uniquely found in B16-F10 samples (J.).

**Figure S6:**
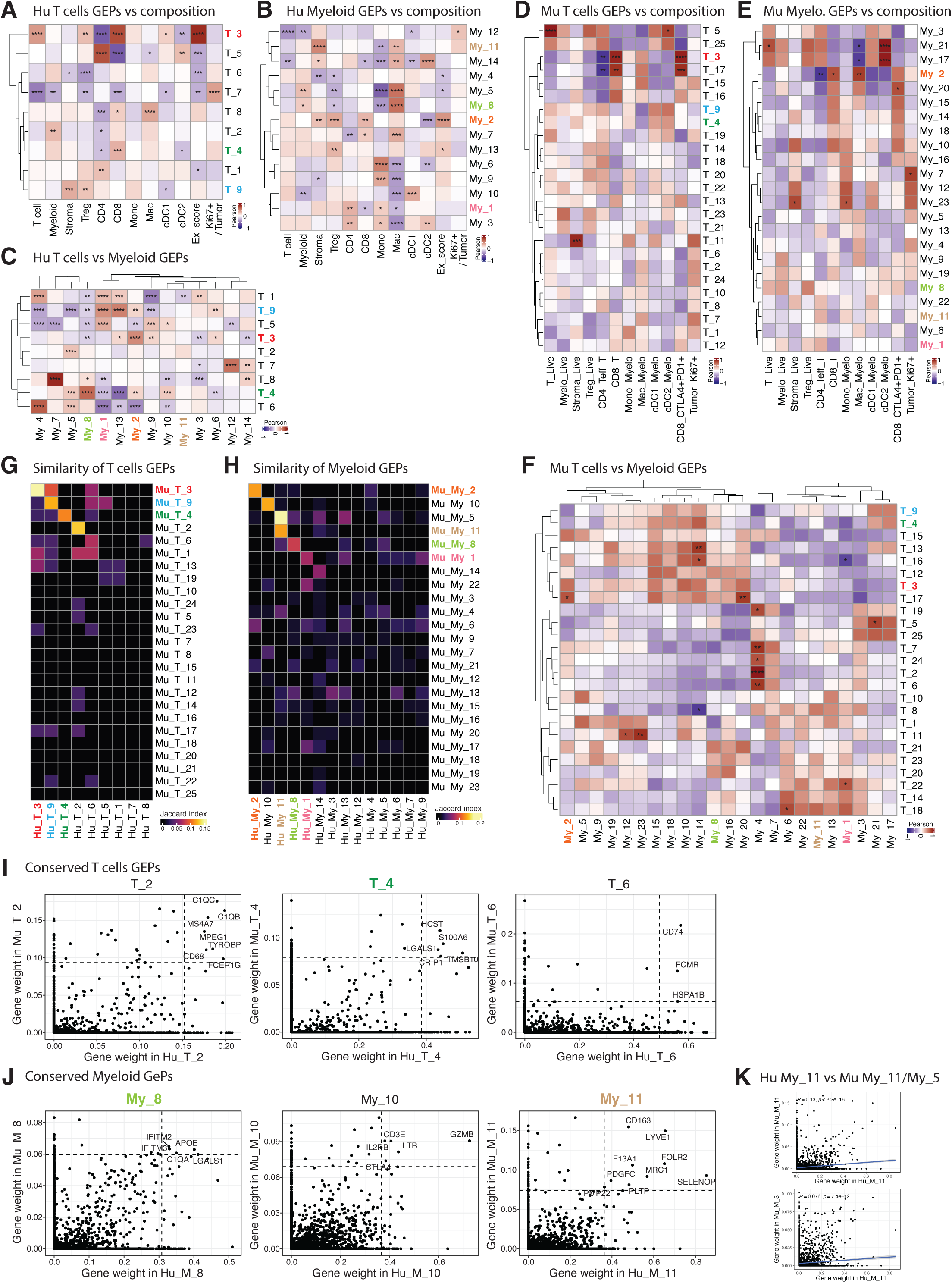
Conservation of human vs murine GEPs, correlations with tumor immune composition and coordinated T vs myeloid GEPs ‘movements’. A. and. **B.** Heatmaps presenting the Pearson correlations of either T cells (A.) or myeloid (B.) GEPs’ enrichment vs the immune composition of human tumor samples. **C.** Heatmaps presenting the Pearson correlations of T cells vs myeloid GEPs’ enrichment across human tumor samples. **D., E. and F.** Heatmaps displaying the same Pearson correlations as in A., B. and C., respectively, across mouse tumor samples. **G. and H.** Heatmaps (non-truncated) showing the Jaccard indexes used to quantify the similarity between human vs mouse GEPs in T cells (G., using top 20 genes per GEP) and myeloid cells (H., using top 50 genes per GEP). Note that although GEPs T_2, T_6 and My_10 appeared similar between mouse and human, we removed them from further analyses as they likely represent a low degree of myeloid contamination/doublets present in T cells for T_2 (Table S1), or conversely for My_10 (Table S1), or show poor overlap of top driver genes for T_6 (Figure S6I and Table S1). **I. and J.** Scatter plots showing the genes contribution (i.e genes weight) to human vs murine conserved T cells (I.) or myeloid (J.) GEPs. Genes annotated show the highest overlapping contributions across species, and the dashed lines separate the 40 highest contributor genes from the others in either factor. **K.** Scatter plots showing the Pearson correlations of the genes weights between human My_11 vs either murine My_11 (left) or murine My_5 (right). Linear regression, R and p values are displayed. Note that both murine My_5 and My_11 show similarity with human My_11 (H.). As these 2 factors seem to be driven by different genes (Figure S5H), we evaluated the correlation of their specific genes’ weights with the ones of Hu_My_11 (K.) to show that Mu_My_11 seem to more closely mimic both the composition (among which LYVE1, FOLR2, CD163, MRC1 and SELENOP) and relative gene weights of Hu_My_11. Statistical significance was calculated using a Pearson correlation test with Benjamini-Hochberg correction, * p.adj ≤ 0.05, ** p.adj ≤ 0.01, *** p.adj ≤ 0.001, **** p.adj ≤ 0.0001.

**Figure S7:**
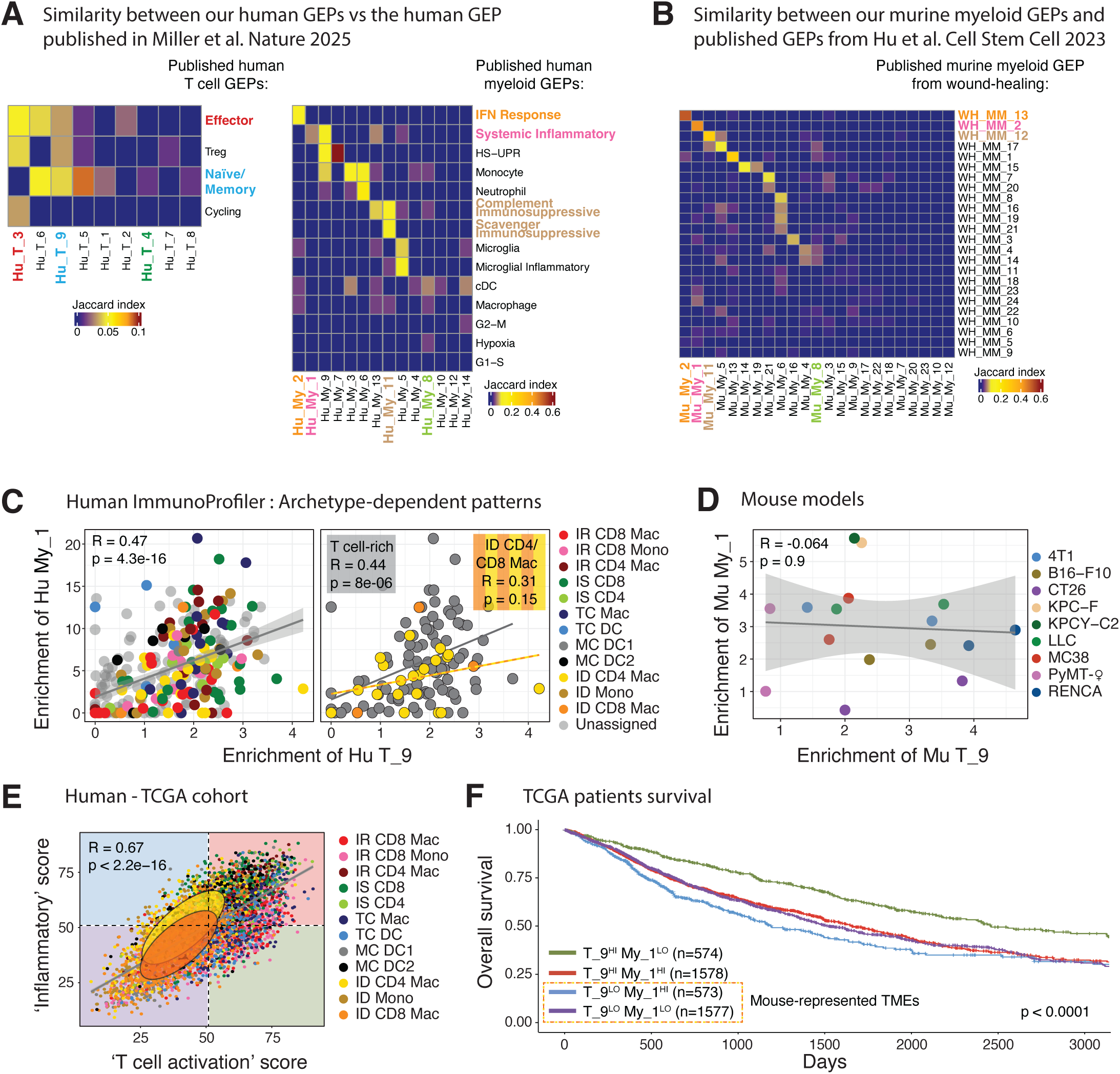
Robustness of our T cells and myeloid GEPs and learning from the discordance of their ‘movements’ between huTMEs vs muTMEs. A. and. **B.** Heatmaps showing Jaccard indexes used to quantify the similarity between (A.) our human T cells (left, using top 20 genes per GEP) and myeloid (right, using top 50 genes per GEP) GEPs to the ones published in^59^ or between (B.) our murine myeloid GEPs and the ones published in^24^ (using top 50 genes per GEP). The top 20 genes for each GEP were used to calculate similarity. **C. and D.** Scatter plots showing the correlation between the enrichments of GEPs T_9 and My_1 found across human (C., colored by archetypes) and mouse tumors (D., colored by tumors lines). Each dot represents a sample, and the diagonal grey lines represent linear regressions. **E.** Scatter plot showing the relative enrichments of gene signatures calculated using the top 20 genes of GEPs T_9 and My_1 in TCGA patients. Patients were binned as High or Low for each GEP if they were present respectively in the top or bottom 50% for each calculated GEP gene score. Each dot represents a sample, and the diagonal grey line represents linear regression. The yellow and orange ellipses in I. highlight the behavior of patients belonging to the Immune Desert CD4/CD8 Macrophages archetypes. **F.** Kaplan-Meier graph showing the overall survival of TCGA patients stratified according to their respective enrichment of T_9 and My_1 following the binning shown in E. Statistical significance in C., D. and E. was calculated using a Pearson correlation test with Benjamini-Hochberg correction. Statistical significance in F. was calculated using a log-rank test.

## References

1. Binnewies, M. et al. Understanding the tumor immune microenvironment (TIME) for effective therapy. Nat Med 24, 541–550 (2018).

2. Bruni, D., Angell, H. K. & Galon, J. The immune contexture and Immunoscore in cancer prognosis and therapeutic efficacy. Nat Rev Cancer 20, 662–680 (2020).

3. Thorsson, V. et al. The Immune Landscape of Cancer. Immunity 48, 812–830.e14 (2018).

4. Bagaev, A. et al. Conserved pan-cancer microenvironment subtypes predict response to immunotherapy. Cancer Cell 39, 845–865.e7 (2021).

5. Pelka, K. et al. Spatially organized multicellular immune hubs in human colorectal cancer. Cell 184, 4734–4752.e20 (2021).

6. Luca, B. A. et al. Atlas of clinically distinct cell states and ecosystems across human solid tumors. Cell 184, 5482–5496.e28 (2021).

7. Combes, A. J. et al. Discovering dominant tumor immune archetypes in a pan-cancer census. Cell 185, 184–203.e19 (2022).

8. Im, K., Combes, A. J., Spitzer, M. H., Satpathy, A. T. & Krummel, M. F. Archetypes of checkpoint-responsive immunity. Trends in Immunology 42, 960–974 (2021).

9. Combes, A. J., Samad, B. & Krummel, M. F. Defining and using immune archetypes to classify and treat cancer. Nat Rev Cancer 23, 491–505 (2023).

10. Ireson, C. R., Alavijeh, M. S., Palmer, A. M., Fowler, E. R. & Jones, H. J. The role of mouse tumour models in the discovery and development of anticancer drugs. Br J Cancer 121, 101–108 (2019).

11. Shay, T. et al. Conservation and divergence in the transcriptional programs of the human and mouse immune systems. Proceedings of the National Academy of Sciences 110, 2946–2951 (2013).

12. Bjornson-Hooper, Z. B. et al. A Comprehensive Atlas of Immunological Differences Between Humans, Mice, and Non-Human Primates. Frontiers in Immunology 13, (2022).

13. Medetgul-Ernar, K. & Davis, M. M. Standing on the shoulders of mice. Immunity 55, 1343–1353 (2022).

14. Leach, D. R., Krummel, M. F. & Allison, J. P. Enhancement of Antitumor Immunity by CTLA-4 Blockade. Science 271, 1734–1736 (1996).

15. Hodi, F. S. et al. Improved Survival with Ipilimumab in Patients with Metastatic Melanoma. New England Journal of Medicine 363, 711–723 (2010).

16. Gengenbacher, N., Singhal, M. & Augustin, H. G. Preclinical mouse solid tumour models: status quo, challenges and perspectives. Nat Rev Cancer 17, 751–765 (2017).

17. Seyhan, A. A. Lost in translation: the valley of death across preclinical and clinical divide – identification of problems and overcoming obstacles. Translational Medicine Communications 4, 18 (2019).

18. Mosely, S. I. S. et al. Rational Selection of Syngeneic Preclinical Tumor Models for Immunotherapeutic Drug Discovery. Cancer Immunol Res 5, 29–41 (2017).

19. Yu, J. W. et al. Tumor-immune profiling of murine syngeneic tumor models as a framework to guide mechanistic studies and predict therapy response in distinct tumor microenvironments. PLOS ONE 13, e0206223 (2018).

20. Taylor, M. A. et al. Longitudinal immune characterization of syngeneic tumor models to enable model selection for immune oncology drug discovery. j. immunotherapy cancer 7, 328 (2019).

21. Gutierrez, W. R. et al. Divergent immune landscapes of primary and syngeneic Kras-driven mouse tumor models. Sci Rep 11, 1098 (2021).

22. Carretta, M. et al. Dissecting tumor microenvironment heterogeneity in syngeneic mouse models: insights on cancer-associated fibroblast phenotypes shaped by infiltrating T cells. Front. Immunol. 14, (2024).

23. Zhong, W. et al. Comparison of the molecular and cellular phenotypes of common mouse syngeneic models with human tumors. BMC Genomics 21, 2 (2020).

24. Hu, K. H. et al. Transcriptional space-time mapping identifies concerted immune and stromal cell patterns and gene programs in wound healing and cancer. Cell Stem Cell 30, 885–903.e10 (2023).

25. Wherry, E. J., Blattman, J. N., Murali-Krishna, K., van der Most, R. & Ahmed, R. Viral Persistence Alters CD8 T-Cell Immunodominance and Tissue Distribution and Results in Distinct Stages of Functional Impairment. Journal of Virology 77, 4911–4927 (2003).

26. Richter, K. et al. Macrophage and T Cell Produced IL-10 Promotes Viral Chronicity. PLOS Pathogens 9, e1003735 (2013).

27. Norris, B. A. et al. Chronic but Not Acute Virus Infection Induces Sustained Expansion of Myeloid Suppressor Cell Numbers that Inhibit Viral-Specific T Cell Immunity. Immunity 38, 309–321 (2013).

28. Philip, M. & Schietinger, A. Heterogeneity and fate choice: T cell exhaustion in cancer and chronic infections. Current Opinion in Immunology 58, 98–103 (2019).

29. McLane, L. M., Abdel-Hakeem, M. S. & Wherry, E. J. CD8 T Cell Exhaustion During Chronic Viral Infection and Cancer. Annual Review of Immunology 37, 457–495 (2019).

30. Binnewies, M. et al. Unleashing Type-2 Dendritic Cells to Drive Protective Antitumor CD4+ T Cell Immunity. Cell 177, 556–571.e16 (2019).

31. Kersten, K. et al. Spatiotemporal co-dependency between macrophages and exhausted CD8+ T cells in cancer. Cancer Cell 40, 624–638.e9 (2022).

32. Suthen, S. et al. Hypoxia-driven immunosuppression by Treg and type-2 conventional dendritic cells in HCC. Hepatology 76, 1329 (2022).

33. Waibl Polania, J., et al. Antigen presentation by tumor-associated macrophages drives T cells from a progenitor exhaustion state to terminal exhaustion. Immunity 58, 232–246.e6 (2025).

34. Fridman, W. H. et al. B cells and tertiary lymphoid structures as determinants of tumour immune contexture and clinical outcome. Nat Rev Clin Oncol 19, 441–457 (2022).

35. Gu-Trantien, C., et al. CXCL13-producing TFH cells link immune suppression and adaptive memory in human breast cancer. JCI Insight 2, (2017).

36. Rodriguez, A. B. et al. Immune mechanisms orchestrate tertiary lymphoid structures in tumors via cancer-associated fibroblasts. Cell Reports 36, 109422 (2021).

37. Ukita, M., et al. CXCL13-producing CD4+ T cells accumulate in the early phase of tertiary lymphoid structures in ovarian cancer. JCI Insight 7, (2022).

38. Jiang, H. et al. Activating Immune Recognition in Pancreatic Ductal Adenocarcinoma via Autophagy Inhibition, MEK Blockade, and CD40 Agonism. Gastroenterology 162, 590–603.e14 (2022).

39. Krishnamurty, A. T. et al. LRRC15+ myofibroblasts dictate the stromal setpoint to suppress tumour immunity. Nature 611, 148–154 (2022).

40. Tharp, K. M. et al. Tumor-associated macrophages restrict CD8+ T cell function through collagen deposition and metabolic reprogramming of the breast cancer microenvironment. Nat Cancer 5, 1045–1062 (2024).

41. Weinstein, J. N. et al. The Cancer Genome Atlas Pan-Cancer analysis project. Nat Genet 45, 1113–1120 (2013).

42. Beura, L. K. et al. Normalizing the environment recapitulates adult human immune traits in laboratory mice. Nature 532, 512–516 (2016).

43. Mempel, T. R., Lill, J. K. & Altenburger, L. M. How chemokines organize the tumour microenvironment. Nat Rev Cancer 24, 28–50 (2024).

44. Kohli, K., Pillarisetty, V. G. & Kim, T. S. Key chemokines direct migration of immune cells in solid tumors. Cancer Gene Ther 29, 10–21 (2022).

45. Groom, J. R. & Luster, A. D. CXCR3 in T cell function. Experimental Cell Research 317, 620–631 (2011).

46. Chow, M. T. et al. Intratumoral Activity of the CXCR3 Chemokine System Is Required for the Efficacy of Anti-PD-1 Therapy. Immunity 50, 1498–1512.e5 (2019).

47. Voabil, P. et al. An ex vivo tumor fragment platform to dissect response to PD-1 blockade in cancer. Nat Med 27, 1250–1261 (2021).

48. Yang, M. et al. CXCL13 shapes immunoactive tumor microenvironment and enhances the efficacy of PD-1 checkpoint blockade in high-grade serous ovarian cancer. J Immunother Cancer 9, e001136 (2021).

49. Zhang, Y. et al. Single-cell analyses reveal key immune cell subsets associated with response to PD-L1 blockade in triple-negative breast cancer. Cancer Cell 39, 1578–1593.e8 (2021).

50. Sorin, M. et al. Single-cell spatial landscape of immunotherapy response reveals mechanisms of CXCL13 enhanced antitumor immunity. J Immunother Cancer 11, e005545 (2023).

51. Magen, A. et al. Intratumoral dendritic cell–CD4+ T helper cell niches enable CD8+ T cell differentiation following PD-1 blockade in hepatocellular carcinoma. Nat Med 29, 1389–1399 (2023).

52. Spranger, S., Dai, D., Horton, B. & Gajewski, T. F. Tumor-Residing Batf3 Dendritic Cells Are Required for Effector T Cell Trafficking and Adoptive T Cell Therapy. Cancer Cell 31, 711–723.e4 (2017).

53. Barry, K. C. et al. A natural killer–dendritic cell axis defines checkpoint therapy– responsive tumor microenvironments. Nat Med 24, 1178–1191 (2018).

54. Broz, M. L. et al. Dissecting the Tumor Myeloid Compartment Reveals Rare Activating Antigen-Presenting Cells Critical for T Cell Immunity. Cancer Cell 26, 638–652 (2014).

55. Stein-O’Brien, G. L. et al. Enter the Matrix: Factorization Uncovers Knowledge from Omics. Trends in Genetics 34, 790–805 (2018).

56. Kinker, G. S. et al. Pan-cancer single-cell RNA-seq identifies recurring programs of cellular heterogeneity. Nat Genet 52, 1208–1218 (2020).

57. Murrow, L. M. et al. Mapping hormone-regulated cell-cell interaction networks in the human breast at single-cell resolution. Cell Systems 13, 644–664.e8 (2022).

58. Gavish, A. et al. Hallmarks of transcriptional intratumour heterogeneity across a thousand tumours. Nature 618, 598–606 (2023).

59. Miller, T. E. et al. Programs, origins and immunomodulatory functions of myeloid cells in glioma. Nature 1–11 (2025) doi:10.1038/s41586-025-08633-8.

60. Benci, J. L. et al. Tumor Interferon Signaling Regulates a Multigenic Resistance Program to Immune Checkpoint Blockade. Cell 167, 1540–1554.e12 (2016).

61. Benci, J. L. et al. Opposing Functions of Interferon Coordinate Adaptive and Innate Immune Responses to Cancer Immune Checkpoint Blockade. Cell 178, 933–948.e14 (2019).

62. Hänggi, K. et al. Interleukin-1α release during necrotic-like cell death generates myeloid-driven immunosuppression that restricts anti-tumor immunity. Cancer Cell 42, 2015–2031.e11 (2024).

63. Caronni, N., Terza, F. L., Frosio, L. & Ostuni, R. IL-1β+ macrophages and the control of pathogenic inflammation in cancer. Trends in Immunology 0, (2025).

64. Remnant, L., Kochanova, N. Y., Reid, C., Cisneros-Soberanis, F. & Earnshaw, W. C. The intrinsically disorderly story of Ki-67. Open Biology 11, 210120 (2021).

65. Salvagno, C. et al. Therapeutic targeting of macrophages enhances chemotherapy efficacy by unleashing type I interferon response. Nat Cell Biol 21, 511–521 (2019).

66. Elewaut, A. et al. Cancer cells impair monocyte-mediated T cell stimulation to evade immunity. Nature 1–10 (2024) doi:10.1038/s41586-024-08257-4.

67. Ray, A. et al. Targeting CD206+ macrophages disrupts the establishment of a key antitumor immune axis. Journal of Experimental Medicine 222, e20240957 (2024).

68. Abolins, S. et al. The comparative immunology of wild and laboratory mice, Mus musculus domesticus. Nat Commun 8, 14811 (2017).

69. Rosshart, S. P. et al. Laboratory mice born to wild mice have natural microbiota and model human immune responses. Science 365, (2019).

70. McGranahan, N. et al. Clonal neoantigens elicit T cell immunoreactivity and sensitivity to immune checkpoint blockade. Science 351, 1463–1469 (2016).

71. Sade-Feldman, M. et al. Defining T Cell States Associated with Response to Checkpoint Immunotherapy in Melanoma. Cell 175, 998–1013.e20 (2018).

72. Friedrich, M. J. et al. The pre-existing T cell landscape determines the response to bispecific T cell engagers in multiple myeloma patients. Cancer Cell 41, 711–725.e6 (2023).

73. Gilbertson, S. E. & Weinmann, A. S. Conservation and divergence in gene regulation between mouse and human immune cells deserves equal emphasis. Trends in Immunology 42, 1077–1087 (2021).

74. Hagai, T. et al. Gene expression variability across cells and species shapes innate immunity. Nature 563, 197–202 (2018).

75. Taneja, P. et al. MMTV mouse models and the diagnostic values of MMTV-like sequences in human breast cancer. Expert Review of Molecular Diagnostics 9, 423–440 (2009).

76. Allen, B. M. et al. Systemic dysfunction and plasticity of the immune macroenvironment in cancer models. Nat Med 26, 1125–1134 (2020).

77. Baron, J. A. et al. The DO-KB Knowledgebase: a 20-year journey developing the disease open science ecosystem. Nucleic Acids Research 52, D1305–D1314 (2024).

78. Bernstein, M. N., Doan, A. & Dewey, C. N. MetaSRA: normalized human sample-specific metadata for the Sequence Read Archive. Bioinformatics 33, 2914–2923 (2017).

79. Wilks, C. et al. recount3: summaries and queries for large-scale RNA-seq expression and splicing. Genome Biology 22, 323 (2021).

80. Gibbs, D. L. Robust classification of Immune Subtypes in Cancer. 2020.01.17.910950 Preprint at 10.1101/2020.01.17.910950 (2020).

81. Ray, A. et al. Multimodal delineation of a layer of effector function among exhausted CD8 T cells in tumors. 2023.09.26.559470 Preprint at 10.1101/2023.09.26.559470 (2025).

82. Brunet, J.-P., Tamayo, P., Golub, T. R. & Mesirov, J. P. Metagenes and molecular pattern discovery using matrix factorization. Proceedings of the National Academy of Sciences 101, 4164–4169 (2004).

83. Pedregosa, F. et al. Scikit-learn: Machine Learning in Python. J. Mach. Learn. Res. 12, 2825–2830 (2011).

